# Existing actin filaments orient new filament growth to provide structural memory of filament alignment during cytokinesis

**DOI:** 10.1101/2020.04.13.039586

**Authors:** Younan Li, Edwin Munro

## Abstract

During cytokinesis, animal cells rapidly remodel the equatorial cortex to build an aligned array of actin filaments called the contractile ring. Local reorientation of filaments by equatorial contraction is thought to underlie the emergence of filament alignment during ring assembly. Here, combining single molecule analysis and modeling in one-cell *C. elegans* embryos, we show that filaments turnover is far too fast for reorientation of single filaments by equatorial contraction/cortex compression to explain the observed alignment, even if favorably oriented filaments are selectively stabilized. Instead, by tracking single Formin/CYK-1::GFP speckles to monitor local filament assembly, we identify a mechanism that we call filament-guided filament assembly (FGFA), in which existing filaments serve as templates to guide/orient the growth of new filaments. We show that FGFA sharply increases the effective lifetime of filament orientation, providing structural memory that allows slow equatorial contraction to build and maintain highly aligned filament arrays, despite rapid turnover of individual filaments.

## Introduction

Non-muscle cells assemble contractile actomyosin arrays to do a variety of jobs, such as cell polarization, cell division, cell migration, wound healing and multicellular tissue morphogenesis (reviewed in (Agarwal and Zaidel-Bar, 2019; Levayer and Lecuit, 2012; Munjal and Lecuit, 2014)). Contractile arrays are assembled from actin filaments, crosslinking proteins, and bipolar myosin II minifilaments, together with various accessory factors that regulate filament assembly, disassembly and motor activity. How assembly/disassembly and activity of contractile arrays are tuned in different ways to build and maintain specific functional architectures remains a fundamental question in cell biology.

One quintessential example of a contractile actomyosin array is the contractile ring, a dynamic network of cross-linked actin filaments and myosin motors that assembles at the cell equator and constricts to divide a single cell into two daughters (Fededa and Gerlich, 2012; Glotzer, 2017; Green et al., 2012; Pollard and O’Shaughnessy, 2019). In animal cells, spatial and temporal control of contractile ring assembly is mediated by equatorial activation of the small GTPase RhoA (Bement et al., 2005; Nishimura and Yonemura, 2006; Yüce et al., 2005). At anaphase onset, local positive and negative signals from the mitotic apparatus specify an equatorial zone of RhoA activity (Glotzer, 2017). RhoA in turn acts through multiple effectors to promote contractile ring assembly. In particular, RhoA binds and activates diaphanous-related formins to promote the local assembly of unbranched actin filaments (Castrillon and Wasserman, 1994; Davies et al., 2014; Grosshans et al., 2005; Watanabe et al., 1997; Watanabe et al., 2010), and it binds and activates Rho Kinase (ROCK) to promote the local recruitment/activation of myosin II (Matsumura, 2005). An initially disorganized network of filaments and motors is then reorganized over time into a more circumferentially aligned array. Although oriented filaments are not required for network contractility, increased filament alignment is associated with increased circumferential tension (Bidone et al., 2014; Spira et al., 2017; Stachowiak et al., 2014) and thus may be important for timely progression and completion of cytokinesis. But how cells build and maintain filament alignment during contractile ring assembly and constriction remains poorly understood.

One model for contractile ring assembly has emerged from intensive studies in fission yeast (Pollard and Wu, 2010). In fission yeast, the contractile ring forms through the coalescence of a broad equatorial band of membrane-attached nodes containing the Anillin-related protein Mid-1 (Wu et al., 2003; 2006). During contractile ring assembly, Mid1p nodes recruit and scaffold the Formin cdc-12 and the Myosin II myp2; cdc-12 promotes the nucleation and elongation of actin filaments, and chance capture of these filaments by myp2 motors on neighboring nodes produces forces that pull nodes together. Cofilin-mediated filament disassembly allows for rapid cycles of search, capture, pull and release (SCPR) (Chen and Pollard, 2011). Empirically constrained computer simulations have shown that SCPR is sufficient to explain the rapid coalescence of nodes from a broad equatorial band into a tight equatorial ring {Vavylonis:2008ho}. The same interactions could also contribute to maintaining filament alignment within the ring, if filament turnover is sufficiently slow (Stachowiak et al., 2014).

In animal cells, the contractile ring forms within a dense array of filaments and motors, in which discrete nodes that anchor sites of filament assembly and/or myosin activity cannot be detected. Borisy and White proposed an alternative to the SCPR model to explain how a dense aligned array of filaments could form at the equator during cytokinesis (White and Borisy, 1983). This model views the cortex as a contractile material, in which local activation of contractility at the equator, and inhibition (relaxation) at the poles, creates a gradient of cortical tension, resulting in a flow of cortical filaments away from the poles and towards the equator. Borisy and White postulated that local realignment of filaments within this flow would lead to buildup of aligned filaments in regions of compressive flow. Compressive flows of cortical material, including actin filaments have been documented during cytokinesis in a variety of different cell types (Fishkind et al., 1996; Khaliullin et al., 2018; Murthy and Wadsworth, 2005; Reymann et al., 2016; Zhou and Wang, 2008). Mathematical models have shown that the realignment of actin filaments by compressive flows could be sufficient to explain the observed degree of filament alignment during contractile ring assembly in some cells (Reymann et al., 2016; Salbreux et al., 2009; White and Borisy, 1983). A key assumption of such models is that individual filaments must be sufficiently stable for flow to build significant alignment within a population of filaments. However, this assumption has yet to be tested by direct simultaneous measurements of flow and filament turnover in any animal cell. Thus, it remains unclear to what extent the local realignment of filaments by cortical flow can explain the rapid emergence and stable maintenance of filament alignment during cytokinesis, and whether additional mechanisms must be involved.

Here we combine TIRF microscopy with single molecule imaging/particle tracking to directly and simultaneously measure rates of contractile flow and filament disassembly during cytokinesis in one-cell *C. elegans* embryos. We find that filament disassembly is far too fast for the alignment of individual filaments by contractile flow to explain the rapid emergence and stable maintenance of filament alignment within the contractile ring. We identify an additional mechanism, in which actin filaments assembled by the Formin CYK-1 use existing filaments as templates to orient their elongation. Combining quantitative image analysis with mathematical modeling, we show that filament-guided filament assembly endows the contractile ring with a structural memory of filament alignment that allows the *C. elegans* embryo to build and maintain a high degree of filament alignment within the contractile ring despite very rapid filament turnover. We propose that similar mechanisms may underlie the assembly and maintenance of aligned filament arrays in many other contexts.

## Results

### Myosin-dependent contractile flow drives the rapid emergence of equatorial filament alignment during contractile ring assembly

To optimize imaging conditions for quantitative analysis of cortical dynamics, we mounted embryos under coverslips using 16µM diameter beads as fixed-size spacers to achieve a uniform degree of compression. Because mild compression can affect overall cortical dynamics (Singh et al., 2019), we began by characterizing the dynamics of contractile ring assembly and furrow ingression in mildly compressed embryos, using near-TIRF microscopy to image probes for cortical F-actin and Myosin II (GFP::UTR (Tse et al., 2012), and NMY-2::mKate2 (Dickinson et al., 2017), Figure 1A, Movie S1, Experimental Procedures). We set the zero timepoint to be an estimate of anaphase onset inferred from the rapid accumulation of equatorial Myosin II ((Werner et al., 2007), Figure 1B). To quantify furrow initiation and progression, we defined equatorial width to be the width of the region that is in focus at the site of furrow ingression, measured perpendicular to the AP axis (Figure 1B). Using measurements of equatorial width and equatorial densities of F-actin and myosin II, we divided early cytokinesis into three phases (Figure 1A, B). Phase I (cortical assembly) begins with anaphase onset and is characterized by rapid accumulation of F-actin and Myosin II with minimal equatorial deformation. Phase II (ring formation/furrow initiation) is characterized by the emergence of filament alignment at roughly constant F-actin and Myosin II density, accompanied the gradual formation of a shallow equatorial furrow. Phase III (ring constriction/furrow ingression) is characterized by a rapid decrease in equatorial width.

**Figure 1.**
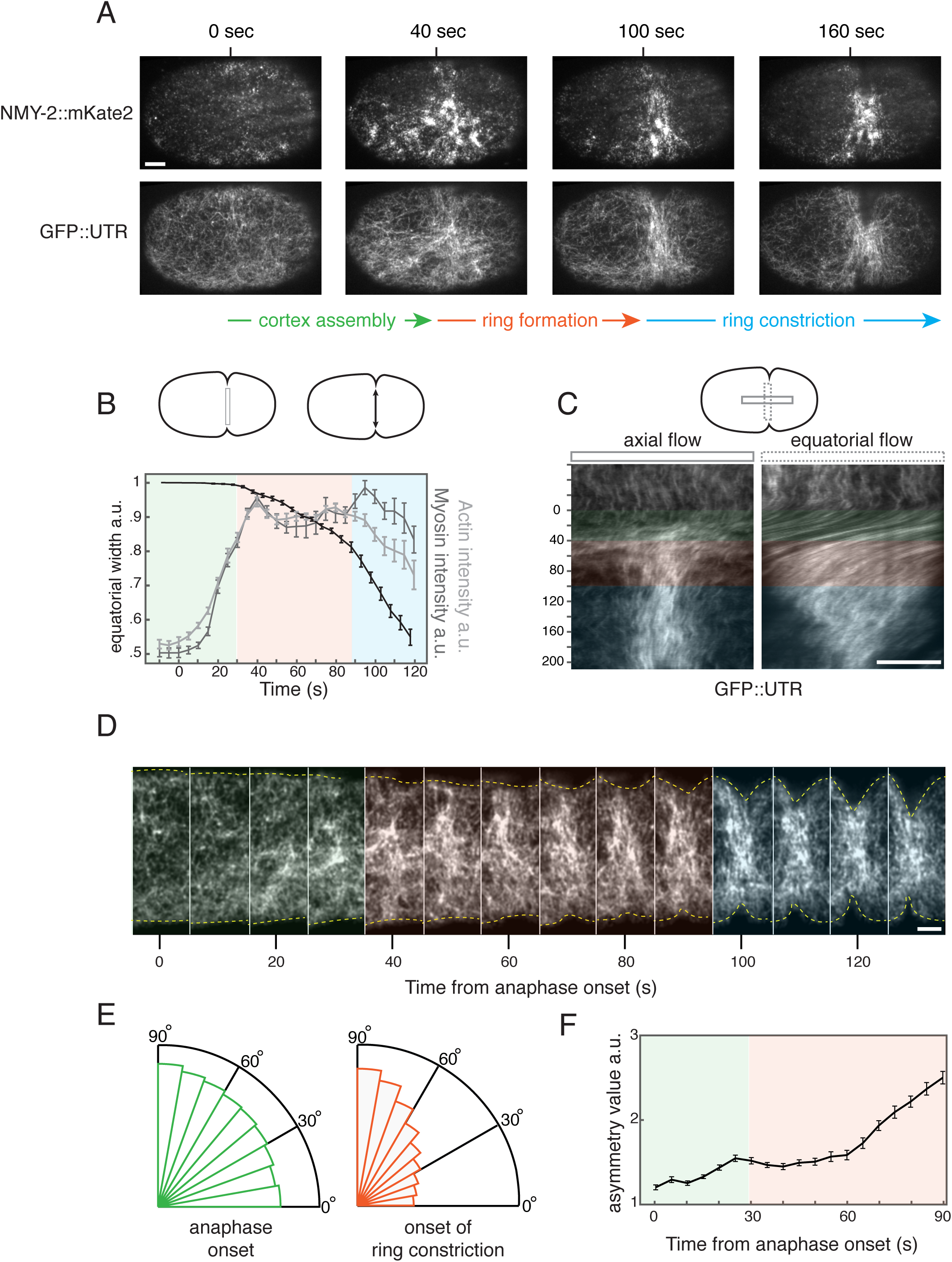
Accumulation and alignment of equatorial actin filaments during cytokinesis. (A) Surface views of cortical myosin II (NMY-2::mKate2, top panels) and F-actin (UTR::GFP, bottom panels) at the indicated time points, measured relative to an estimate of anaphase onset (see text for details). Anterior is to the left in this and all subsequent figures. Scale bars = 5µm. See also Move S1; (B) Measurements of equatorial width (black), mean intensities of F-actin (UTR::GFP, light grey) and myosin II (NMY-2:mKate2, dark grey) over time during cytokinesis. Color overlays indicate the three phases of cytokinesis defined by these measurements: (I) cortex assembly (green), (II) ring formation/furrow initiation (red), and (III) ring constriction (blue). Error bars indicate SEM (n = 5 embryos). Top schematic indicates the regions used for measurements of probe densities (box), and equatorial width (double headed black arrow). (C) Kymographs showing axial (left) and equatorial (rotational) (right) cortical flows for the embryo in (A) and in Movie S1. Top schematic indicates the regions used to make axial (solid box) and equatorial (dashed box) kymographs. Color overlays mark the three phases of cytokinesis. Scale bar = 5µm. (D) Magnified views of UTR::GFP on the equatorial cortex, from the embryo in (A), showing the emergence of filament alignment over time. Yellow dashed lines indicate the cell boundary. Color overlays mark the three phases of cytokinesis. Scale bar = 2µm. (E) Radial histograms showing the distribution of local filament orientations at the equator at anaphase onset (left) and at the onset of ring constriction (right) (n = 5 embryos). (F) Plot of mean asymmetry value vs. time measured on the equatorial cortex (6 µm in width) during early cytokinesis. Error bars indicate SEM (n = 5 embryos).

We used kymography to characterize the pattern and timing of cortical flows that accompany contractile ring assembly in mildly compressed embryos. Axial kymographs revealed a transient posterior to anterior flow that begins with anaphase onset and transitions to a compressive flow, centered on the equator, that persists through phases II and III (Figure 1C; left kymograph).We also observed a transient rapid cortical rotation, perpendicular to the AP axis (Singh et al., 2019) that begins with anaphase onset, and then attenuates during phases II and III (Figure 1C; right kymograph).

To quantify the emergence of equatorial filament alignment during Phases I and II (Figure 1D), we used a standard approach, based on the Sobel operator (Gonzalez and Woods, 2017) to measure the amplitudes and directions of local gradients in the intensity of GFP::UTR, which correspond to individual filaments and/or small bundles (Figure S1A,B, Experimental Procedures). To estimate the distribution of filament orientation, we selected all pixels with gradient amplitudes above a threshold level, and computed a normalized distribution of intensity gradient directions, weighted by gradient amplitude (Figure 1E, Figure S1A,B). Finally, we computed a simple index of filament alignment asymmetry by calculating the ratio of histogram densities within 10° of equatorial (90°) and axial (0°) directions (Figure 1F). Consistent with direct observations, the distribution of equatorial filament orientations was approximately isotropic at anaphase onset, with asymmetry values close to 1.2 (Figure 1E,F). The asymmetry value increased slowly and steadily during Phase I and early Phase II, and then increased more sharply during late Phase II, reaching a mean value of ∼2.5 at the onset of furrow ingression (Figure 1E,F). In contrast, the distribution of polar filament orientations remained largely isotropic from anaphase onset through the end of Phase II. (Fig S1E). Measurements in fixed, phalloidin-stained embryos suggest that equatorial filaments become even more aligned in uncompressed embryos (Figure S1C), but here we used the lower values for direct comparison with the single molecule measurements reported below.

Importantly, in NMY-2:: mKate2; GFP::UTR embryos strongly depleted of NMY-2 by RNAi, a rapid increase in equatorial actin filaments occurred with similar timing after anaphase onset (Figure S2A,B). However, both axial and equatorial cortical flows were essentially abolished (Figure S2C), and the distribution of equatorial actin filament orientations remained largely isotropic after ∼90 seconds, corresponding to the end of Phase II in control embryos (Figure S2D). Thus, in agreement with previous work (Reymann et al., 2016), myosin-driven contraction is required for the rapid emergence of filament alignment prior to ring constriction.

### A simple model reveals the dependence of filament alignment on filament turnover and equatorial contraction rate

The emergence of equatorial filament alignment will depend on three factors: the rate and orientation of local filament assembly; local realignment of existing filaments; and the rate and orientation-dependence of local filament disassembly (Figure 2A). To establish a quantitative framework for assessing how the emergence of filament alignment is shaped by the interplay of these factors, we modeled a population of filaments within a patch of equatorial cortex that undergoes compression at a constant contraction rate *ξ* (Figure 2B). We assumed that filaments assemble within the patch at a rate *k*_*ass*_(*θ*) and disassemble at a rate *k*_*diss*_(*θ*)*ρ*(*θ*), where *ρ*(*θ*) is the density of filaments with orientation *θ*. The patch boundaries move with local flow, such that there is no net movement of filaments into/out of the patch (Figure 2B). Finally, we assumed that the cortical filament network undergoes locally affine deformation, i.e. the change in a filament’s orientation is determined only by the movements of its endpoints, such that the rate of change of a filament’s orientation is given by:

**Figure 2.**
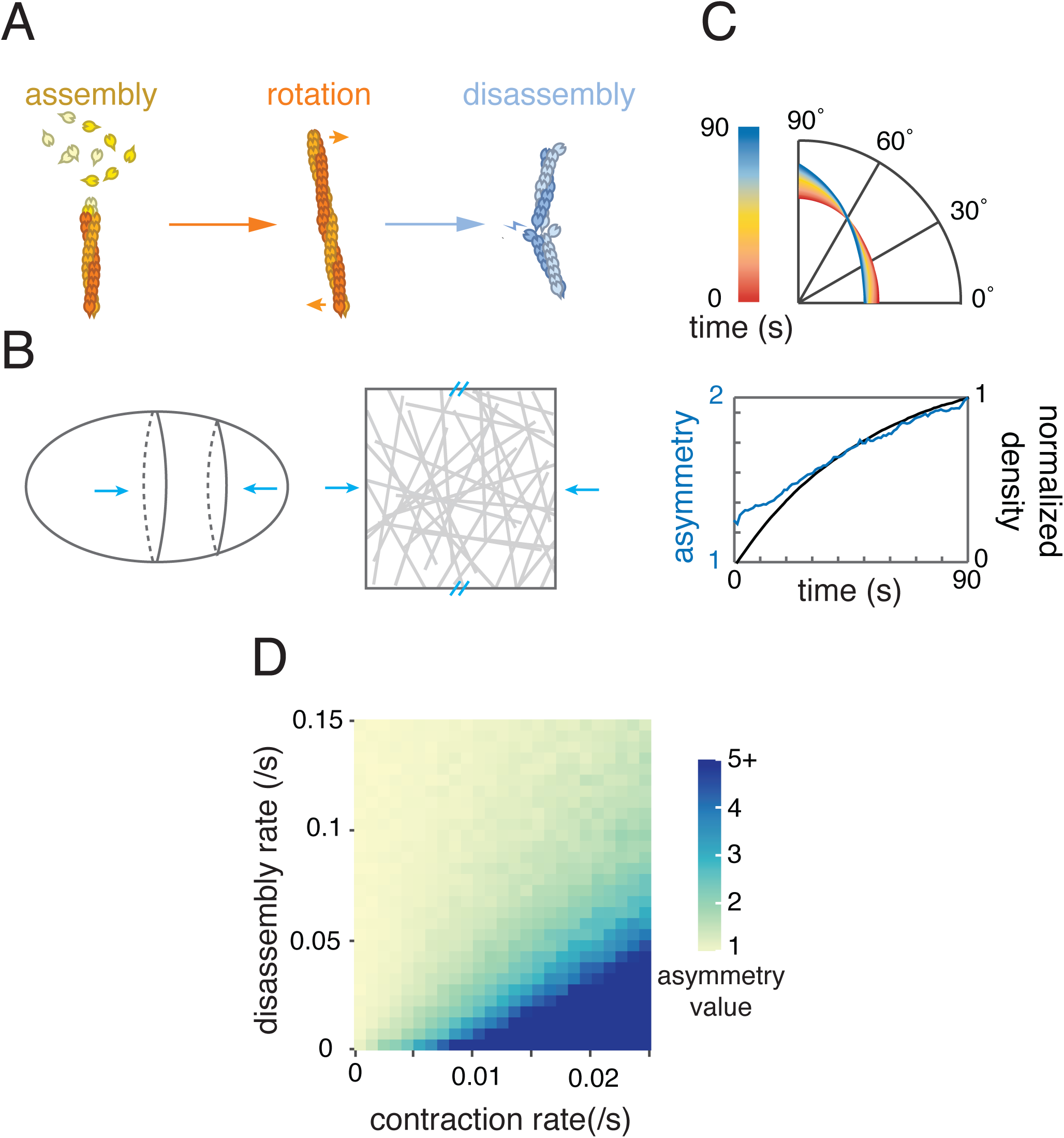
A simple model for filament realignment in compressive flow. (A) Schematic overview of the three processes by which filament number and orientation are assumed to change: local assembly, rotation by contractile flow, and local disassembly. (B) Model representation of a patch of equatorial cortex undergoing uniform axial compression. Top and bottom boundaries of the patch are fixed, while left and right boundaries move at the rate of cortical flow, such that there is no flux of filaments across those boundaries. (C) Model output for one choice of model parameters. Top: Time evolution of the filament orientation distribution. Bright red line indicates the initial distribution, as shown in Figure 1E. Bright blue line indicates the distribution predicted after 90 seconds. Bottom: Plot of the asymmetry value (blue) and normalized filament density vs. time (black). Parameters: contraction rate = 0.01 /s, disassembly rate = 0.03 /s. (D) Simulation outcomes for different values of disassembly and contraction rates, color coded for the asymmetry value achieved after 90 sec.

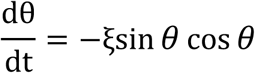

With these assumptions (see Modeling Procedures), we can write an equation that governs the time evolution of filament orientations within the contracting equatorial patch:

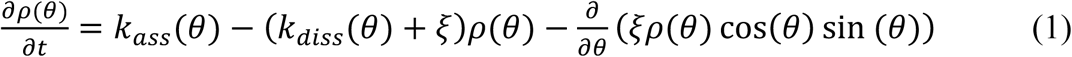

We first considered a simple scenario in which assembly and disassembly rates lack orientation bias or dependence (*k*_*ass*_(*θ*) = *k*_*ass*_; *k*_*diss*_(*θ*) = *k*_*diss*_). For this scenario, given a constant contraction rate, the total density of filaments will approach a steady state level given by 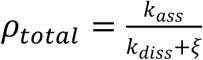 (see Modeling Procedures). Scaling *ρ*(*θ*) by *ρ*_*total*_, we obtained:

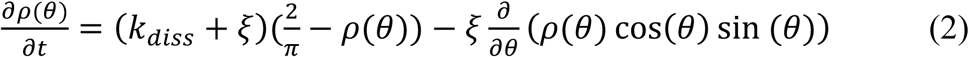

The first term in equation 2 represents the continuous evolution of filament orientations towards a flat (isotropic) distribution, at a characteristic rate *k*_*diss*_ + *ξ*. The second term represents the continuous reorientation of filaments by flow, driving the distribution of filament orientations away from isotropic. Thus, for this simple scenario in which filament assembly /disassembly rates do not depend on filament orientation, the distribution of filament orientations will depend only on the disassembly and contraction rates *k*_*diss*_ and *ξ*. To characterize this dependence, we implemented our model as a simple stochastic simulation (see Experimental Procedures), initialized with the distribution of filament orientations observed during Phase I, and ran for 90 secs, corresponding to the total duration of Phases I and II (Figure 2C). Plotting the asymmetry value after 90 seconds as a function of k_diss_ and *ξ* confirms that asymmetry increases with faster contraction and slower disassembly, and reveals the range of values for which simulations reproduce the observed asymmetries (Figure 2D)

### Filament turnover is too fast for reorientation of actin filaments by cortical flow to explain the emergence of equatorial filament alignment

To test these predictions, we used single molecule imaging and particle tracking as previously described (Robin et al., 2014) to measure contraction and disassembly rates during cytokinesis *in vivo*. Briefly, we used near-TIRF microscopy to image embryos expressing Actin-GFP at single molecule levels (Figure 3A), from anaphase onset through the onset of ring constriction. We performed particle-tracking analysis to obtain a dense sampling of single molecule trajectories throughout the cortex and over time (Figure 3B, Movie S2).

**Figure 3.**
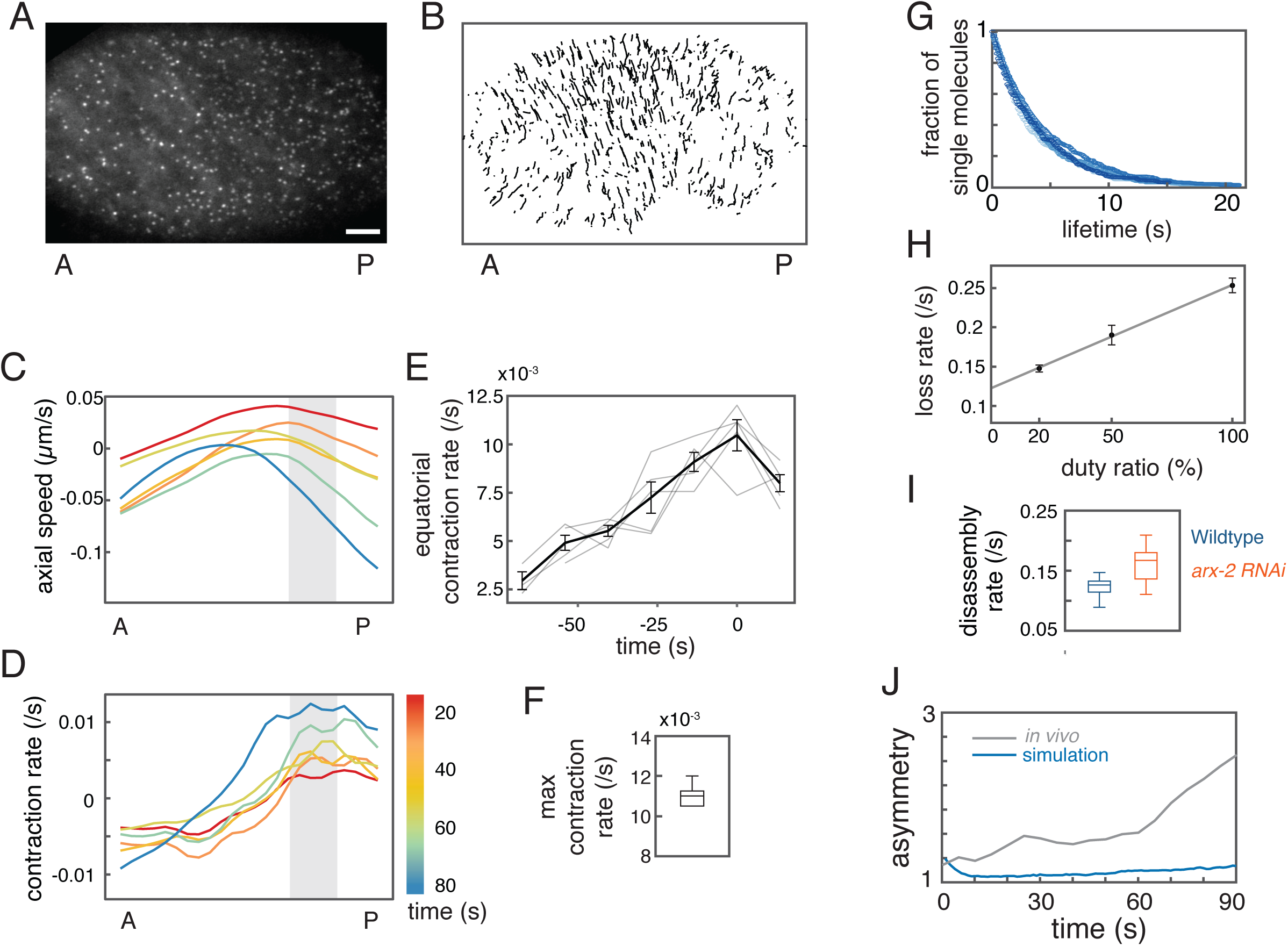
Simultaneous single molecule measurements of contraction and disassembly rates *in vivo*. (A) Near-TIRF image of an embryo expressing Actin::GFP at single molecule levels, taken under standard conditions for single particle tracking. See also movie S2. Scale bar = 5µm. (B) Trajectories of all single molecules tracked over a 22.5 sec time interval. (C) Mean axial velocity of single molecules measured at different positions along the anterior-posterior axis at different time points during cytokinesis for one sample embryo. A negative value indicates anterior movement. (D) Mean contraction rate measured as the spatial derivative of axial speeds shown in (C). Gray boxes in (C&D) indicate the equatorial region (10µm in width), and the color map indicates the time at which each curve was measured relative to anaphase onset. (E) Mean equatorial contraction rate vs. time measured for five different embryos. Plots are aligned relative to the onset of phase III (furrow constriction). Gray lines show data for individual embryos. Black line and error bars indicate the mean +/- SEM (n=5 embryos). (F) Maximum equatorial contraction rate measured just before constriction in wild-type embryos (n=5). (G) Loss curves plotting the fraction of single molecule trajectories with a given lifetime (n=5). (H) Plots of loss rate vs duty ratio. Data points and error bars indicate mean +/- SEM (20%: n = 5 embryos; 50%: n = 5 embryos; 100%: n = 5 embryos). Solid line indicates fit to loss rate = disassembly rate + photobleach rate * duty ratio. (I) Disassembly rate measured in the equatorial region during late phase II (30 seconds prior to phase III) for wild type (n = 5, blue) and *arx-2 (RNAi)* (n = 6, green) embryos. (J) Model prediction of filament asymmetry over time given measured contraction rate (E) and disassembly rate (I). Gray line shows the mean filament asymmetry measured during cytokinesis in vivo (from Figure 1F). Blue line indicates the model-predicted asymmetry value.

To quantify cortical flow, we sampled single molecule displacements over 13.5 second intervals to estimate local actin filament velocities. We then binned these data to produce estimates of mean axial velocity as a function of position and time during Phases I and II (Figure 3D, Experimental Procedures). This analysis confirmed the characteristic pattern of axial cortical flow revealed by kymographs (Figure 1B, left), with an early posterior-anterior flow giving way to compressive flow from both poles towards the equator, with maximum speeds that increased over time (Figure 3D). Notably, we observed only small deviations of single molecule movement from the bulk flow of surrounding molecules, confirming that the cortical network undergoes a locally affine deformation (data not shown). The spatial derivative (the slope) of the axial velocity was approximately constant and negative within an equatorial region about 10 *μm* wide (gray region in Figure 3D), indicating a region of uniform local contraction. Plotting the average equatorial contraction rate as a function of time, using the onset of rapid furrow ingression (Phase III) to align data from multiple embryos, revealed a steady increase in contraction rate from anaphase onset through the onset of rapid furrow ingression, approaching a maximum value of 0.011 +/- 0.001/sec (n = 5 embryos; Figure 3E), in agreement with previous estimates made by other methods (Khaliullin et al., 2018; Reymann et al., 2016).

To extract local estimates of F-actin disassembly rate from single molecule trajectories, we compiled the trajectories collected within a given time window and region of interest, and then constructed standard decay curves by plotting the number of trajectories with length > *τ* sec for different values of *τ* (Figure 3G). These decay curves were well-fit by single exponentials, indicating a single loss rate, *k*_*loss*_, which is the sum of an intrinsic filament disassembly rate *k*_*dis*_ and a photobleaching rate, *k*_*pb*,_ which depends on laser power and duty ratio (i.e. the fraction of time the laser is on). To obtain separate measurements of disassembly and photobleaching rates, we collected a sequence of measurements from different embryos, holding laser power and exposure time constant and varying the duty ratio. These data were well-fit by a function of the form: *k*_*loss*_ = *k*_*dis*_ + *k*_*pb*_ · *dr* (Figure 3H, S3A), yielding an estimate of the actin filament disassembly rate k_diss_ = 0.122/s (Figure 3I), which agrees well with the value we previously reported for embryos depleted of Myosin II (Robin et al., 2014).

Because GFP-tagged actin monomers incorporate less efficiently into formin-assembled filaments in other contexts (Chen et al., 2012), our measurements could reflect a biased contribution from disassembly of branched actin filaments. To address this concern, we measured filament disassembly rates in embryos strongly depleted of the ARP2/3 complex subunit ARX-2. We again observed mono-exponential decay kinetics, with a slight increase in the estimated disassembly rate to k_diss_ = 0.167/s. Thus our approach may slightly underestimate the disassembly rate of formin-assembled filaments within the contractile ring.

Using the measured contraction and disassembly rates, our simple model predicts that filament alignment asymmetry will first decrease from an initial value of 1.2, and then slowly increase to reach a value of 1.19 after 90 seconds (Figure 3J). We concluded that given constant filament turnover, reorientation of individual filaments by cortical flow cannot explain the rapid emergence of filament alignment during cytokinesis in *C. elegans* zygotes.

### Preferential stabilization of correctly oriented filaments cannot explain robust alignment

One possibility is that correctly oriented filaments are preferentially stabilized during ring assembly. For example, local crosslinking or bundling of co-aligned filaments by factors like Anillin could protect them from disassembly (Tian et al., 2015). To ask whether such an effect could explain the rapid emergence of filament alignment in *C. elegans* zygotes, we introduced into our model a generic form of orientation dependence, in which the disassembly rate is governed by a Hill function of filament orientation (Figure S4A):

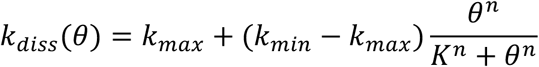

We then performed a series of simulations in which we systematically varied K, n, *k*_*min*_ and *k*_*max*_, using our measured values for contraction rate vs time (Figure 3E). We identified a subset of parameter values for which simulations predicted mean disassembly rates and asymmetry values (after 90 sec) close to those measured *in vivo* (Figure S4B; 2 < asymmetry value < 3; 0.1/s < mean k_diss_ < 0.2/s). Strikingly however, for none of these values could simulations also reproduce the time-dependent changes in filament density and asymmetry measured *in vivo* (Figure S4C,D). *In vivo*, a rapid rise in filament density precedes a stable plateau in Phase II, while a slow rise in asymmetry precedes a rapid rise in late Phase II (Figure S4D). By contrast, in simulations, rapid rises in filament density and asymmetry were invariably correlated (Figure S4E) for a simple reason: if filament stability increases with filament alignment, then a sharp increase in filament asymmetry will inevitably produce a sharp increase in filament density (assuming constant assembly rate). Thus, a rapid increase in filament asymmetry cannot coincide with a stable plateau in filament density, as we observe in late phase II, and thus orientation-dependent filament disassembly cannot alone explain the rapid emergence of filament alignment during contractile ring assembly.

### CYK-1-dependent filament elongation is directionally biased at the equator during cytokinesis

An alternative possibility is that the orientation of actin filament assembly could be biased. To test this possibility, we developed an approach to measure the orientation of filament elongation during cytokinesis in embryos expressing an endogenously-tagged form of the formin CYK-1 (CYK-1::GFP) (Padmanabhan et al., 2017). CYK-1 is required for contractile ring assembly and cytokinesis in early *C. elegans* embryos (Davies et al., 2014; Severson et al., 2002; Swan et al., 1998). Like other diaphanous-related formins, CYK-1 dimers presumably associate with the barbed ends of rapidly elongating actin filaments (Figure 4A). Therefore, directional movements of CYK-1 molecules should provide a direct readout of the orientation of filament growth.

**Figure 4.**
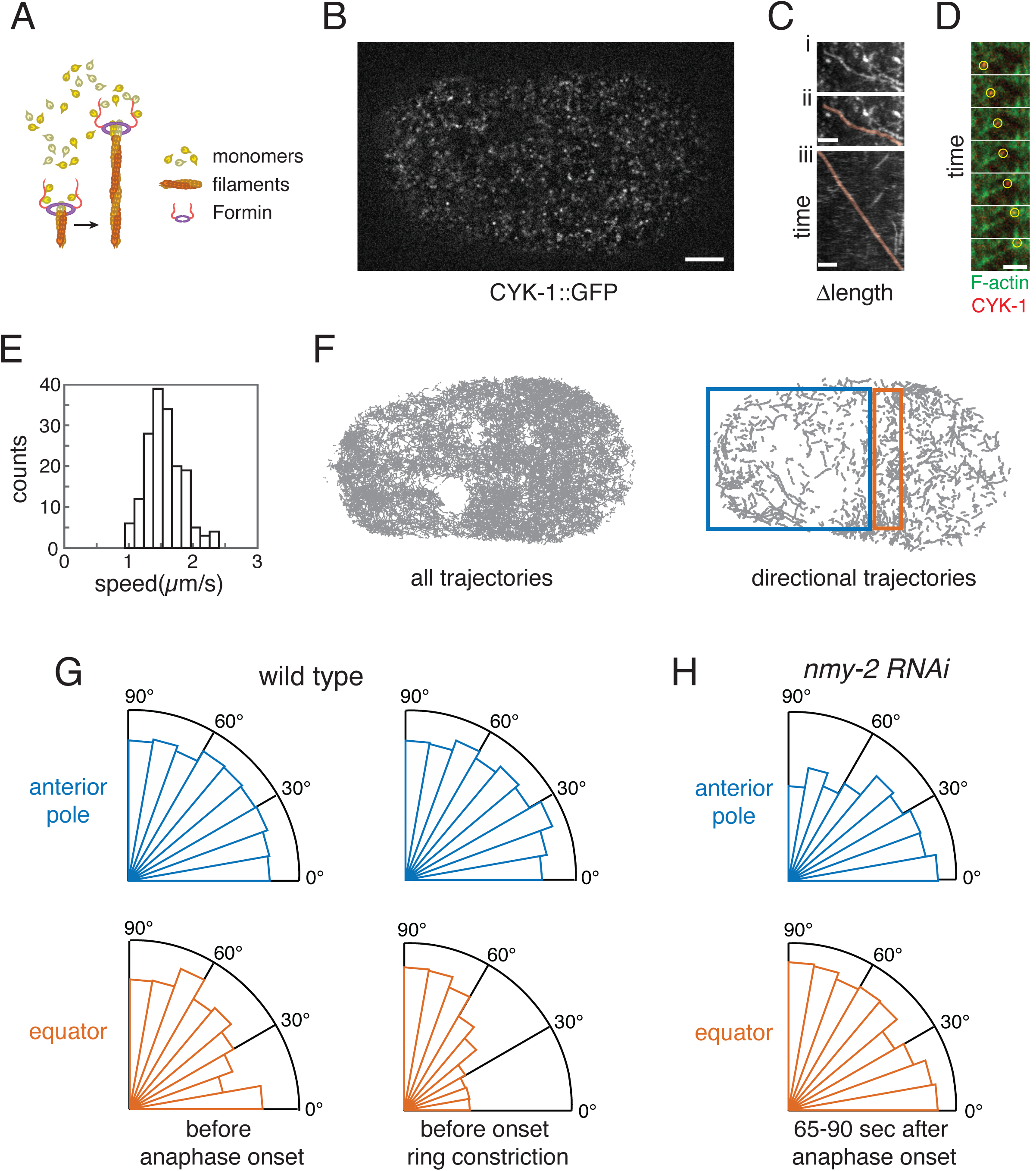
Orientation of Formin-dependent filament elongation is biased with respect to the equatorial axis during cytokinesis. (A) Schematic view of formin-mediated actin filament assembly. Formin dimers remain associated with the barbed ends of elongating filaments. (B) Surface view of a one-cell embryo at anaphase onset expressing CYK-1::GFP, and processed to highlight moving CYK-1::GFP speckles (see Experimental Procedures and Movie S3), Scale bar = 5µm. (C) Focus on a single CYK-1::GFP trajectory during cytokinesis in a wild-type embryo. (i) Maximum intensity projection of a small region of interest over time reveals moving CYK-1 particles. (ii) Trajectory of a single moving CYK-1::GFP particle is highlighted in orange. (iii) Kymograph made by straightening the orange trajectory reveals ∼constant speed. Scale bar = 2µm. (D) Sequence of images from the polar region of an embryo expressing CYK-1::GFP/LifeAct::mCherry and partially depleted of NMY-2. Yellow circles highlight a CYK-1::GFP particle moving into a region of low F-actin density, leaving a newly assembled actin filament behind it. See also Movie S5. Scale bar = 2µm. (E) Distribution of elongation rates measured for 41 rapidly directionally moving CYK-1::GFP speckle trajectories broken into 170 0.6 sec segments (see Experimental Procedures for details). (F) CYK-1 trajectories identified by automated particle tracking analysis. Left panel: all trajectories with lifetime >= 0.6 seconds. Right panel: The subset of trajectories selected for fast directional movement (see Experimental Procedures for details, and Movie S4). Rectangles indicate polar (blue) and equatorial (orange) regions used for measurements shown in (G) and (H). (G) Distribution of movement orientations for the fast directionally moving cortical CYK-1 particles at the equator (orange) and anterior pole (blue) just before anaphase onset (left) and just before the onset of ring constriction (right). Data pooled from n = 5 embryos. (H) Distribution of movement orientations for the fast directionally moving cortical CYK-1 particles in *nmy-2 (RNAi)* embryos at the equator and anterior pole 65-90 seconds after anaphase onset. Data pooled from n = 5 embryos.

Using near-TIRF microscopy, we could detect CYK-1::GFP at the cortex as diffraction-limited speckles (Figure 4B). Many of these speckles are stationary, while the remainder undergo rapid directional movement (Figure 4C, Movie S3). Fast dual-color imaging of CYK-1::GFP and LifeAct::mCherry revealed the rapid appearance of newly assembled filaments behind a subset of fast-moving CYK-1::GFP speckles moving through regions of low F-actin density (Figure 4D, Movie S5), confirming that these speckles mark the barbed ends of actively elongating actin filaments. If immobile CYK-1::GFP speckles are engaged in elongating filaments, those filaments should move rapidly away from stationary CYK-1::GFP speckles as they elongate. However, using particle tracking analysis of single-molecule speckles of Actin::GFP or UTR::GFP, we could not detect a pool of rapidly-moving GFP speckles (Figure 3B and data not shown). Thus it is likely that the stationary CYK-1::GFP speckles represent inactive protein, while the fast moving CYK-1::GFP speckles represent CYK-1 dimers associated with the barbed ends of rapidly elongating filaments.

To characterize the orientation of filament growth, we developed methods to detect, track and analyze fast, directionally moving CYK-1 speckles (Movie S4, See Experimental Procedures for details). Briefly, we subtracted a moving minimum intensity projection (∼800 msec window) from the raw data to enhance the signal associated with moving CYK-1 speckles. We performed particle detection and tracking on this filtered data. We first hand-picked and verified 41 fast moving CYK-1 trajectories, subdivided them into shorter (10 frame = 0.6 second) segments, and calculated their average instantaneous speed to be 1.561 +/-0.286 um/s (Figure 4E), which is similar to the elongation speeds previously measured for the formin mDia2 in XTC fibroblasts (Higashida et al., 2004). Then we subdivided all trajectories into ten-frame segments, and used a statistical filter developed by Jaqaman et. al. (2008) to select the subset representing directional motion, whose mean velocities were within the range measured for the hand-picked trajectories in Figure 4E (Figure 4F). Finally, we fit a straight line to each segment to estimate elongation direction as the angle between the fitted line and the embryo’s AP axis.

We then plotted the distribution of elongation directions in equatorial and polar regions during 24-second windows of time just before anaphase onset, and just before the onset of rapid furrow ingression (Figure 4G). Before anaphase onset, filament elongation was approximately isotropic in both equatorial and polar regions (Figure 4G; asymmetry values = 0.96 at the equator and 0.99 at the poles. equator: n = 1380 trajectory segments; pole: n = 2978 trajectory segments in 5 embryos). Just before the onset of furrow ingression, elongation within the polar region remained isotropic, however, in the equatorial region, elongation was strongly biased perpendicular to the AP axis (Figure 4G; asymmetry values = 2.19 at the equator and 1.03 at the poles. equator: n = 1469 trajectory segments; pole: n = 3029 trajectory segments in 6 embryos). Thus, biased orientation of actin filament assembly underlies the emergence of filament alignment during contractile ring assembly.

### Newly assembled filaments use existing filaments to orient their elongation

These results reveal a correlation between the orientations of existing filaments and the elongation of new filaments: Actin filament orientation and growth are both isotropic on the polar cortex throughout cytokinesis, and at the equator before anaphase onset (Figure 1E &S1E, Figure 4G). Filament orientation and growth are anisotropic and co-aligned at the equatorial cortex before ring constriction (Figure 1E and Figure 4G). Importantly, in embryos strongly depleted of NMY-2 by RNAi, equatorial filaments remain isotropic 65-90 seconds after anaphase onset (Figure S2A,E), and equatorial filament assembly was also isotropic (Figure 4H; asymmetry value = 0.99; n = 2099 trajectory segments in 4 embryos). These results rule out the possibility that some other equatorial signal biases filament elongation independent of existing filament’s orientations. Instead, they suggest that aligned equatorial filaments provide a local directional cue to orient new filament assembly.

To examine this further, we analyzed the behavior of CYK-1::GFP speckles in embryos expressing both CYK-1::GFP and LifeAct::mCherry. In polar regions, where the density of filaments is lower, a significant fraction of fast-moving CYK-1 speckles moved along existing actin filaments or small filament bundles. Moreover, in many cases, moving CYK-1 speckles altered direction upon encountering an existing filament/bundle, and then subsequently moved along that filament/bundle (Figure 5A, Movie S6). These observations strongly suggest that existing filaments provide cues that act locally and continuously to bias the direction of new filament elongation.

**Figure 5.**
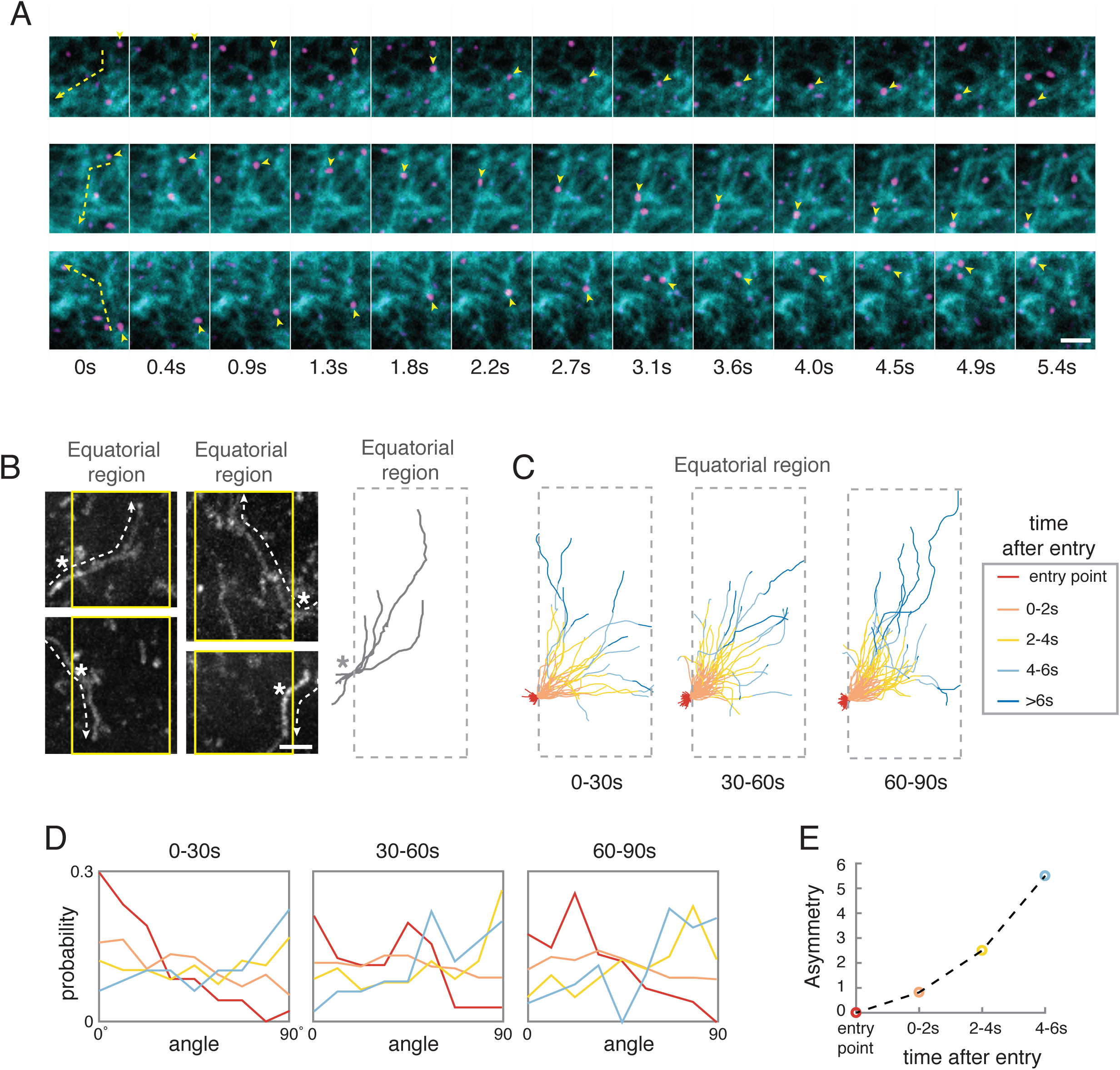
Elongating filaments use existing filaments to orient their growth: (A) Image montages highlighting three different filament growth trajectories in a one-cell embryo expressing transgenic CYK-1::GFP/LifeAct::mCherry and partially depleted of NMY-2. Yellow arrow heads mark the position and direction of a rapidly moving CYK-1::GFP speckle; dashed yellow lines and arrows at the left mark actin filament/bundles that the CYK-1 speckle moves along. In all examples CYK-1 speckles move along one actin filament/bundle and then change direction as they encounter a second filament/bundle. See also Movie S6. Scale bars = 2µm. (B) Left panel: Four example CYK-1 trajectories entering the equatorial region from the left or right and turning either up or down to move along the equatorial axis. Each image is a maximum intensity projection of many frames. Dashed lines and white arrows mark the individual CYK-1 trajectories; white asterisks mark the point of entry into the equatorial region, and vertical yellow lines indicate the boundary of the equatorial region (6um in width, see also Movie S7). Right panel: For quantitative analysis in (C) and (D), trajectories were aligned with respect to their equatorial entry point, and flipped vertically and/or horizontally so that all enter from the left and turn upwards. Scale bars = 2µm. (C) Superposition of many CYK-1 trajectories entering the equatorial region (6*μm* in width) in different time windows, measured relative to anaphase onset. Individual trajectory segments are color-coded based on the time after entry into the equatorial region (see legend at right). (D) Probability histograms showing distributions of CYK-1 movement orientations at different times after entry into the equatorial region, in different time windows during cytokinesis (showing on top of histogram). Individual histograms are color-coded as in (C). (E) Line plot showing the asymmetry value of CYK-1 movement orientations at different times after entry into the equatorial region during the 60-90 second time window (LATE PHASE II).

To quantify the strength of this effect at the equator, we analyzed the behavior of 202 CYK-1::GFP speckles from 13 embryos that began outside, and moved into, the equatorial region (Figure 5B, Movie S7). We aligned all 202 trajectories with respect to their point of entry into the equatorial region and reflected trajectories about the AP and/or equatorial axis to place them all into the same quadrant (Figure 5B). We grouped whole trajectories based on time after anaphase onset (0-30sec, 30-60sec, 60-90sec), and then analyzed changes in individual trajectory orientations based on time after entry into the equatorial region (Figure 5C-D). We observed a shift in trajectory orientations within all three windows of time (Figure 5D). However, the most dramatic shift occurred just before onset of ring constriction (60-90sec), when equatorial filaments are highly aligned (Figure 5D). In this window, the mean asymmetry of filament growth increased from a value of ∼ 0 at the time of entry into the equatorial region to ∼ 6 five seconds later (Figure 5E). We conclude that aligned equatorial filaments provide a strong local directional cue to orient the elongation of newly assembling filaments. Hereafter we refer to this biased elongation as filament-guided filament assembly (FGFA).

### Filament-guided filament assembly (FGFA) increases the “effective lifetime” of filament orientation, and can explain the observed degree of filament alignment, given measured contraction and disassembly rates

To further characterize how FGFA shapes the emergence of filament alignment, we modified our original model to incorporate a simple representation of this orientation cue. We assumed that a fraction *λ* of growing filaments elongate in the direction of an existing filament (i.e. with an orientation chosen at random from the distribution of existing filament orientations), while the remainder (1-*λ*) elongate with random orientations (Figure 6A). With these assumptions, and scaling filament density so that the total filament density approaches a value of 1 (see modeling procedures), we obtain a modified version of equation (2):

**Figure 6.**
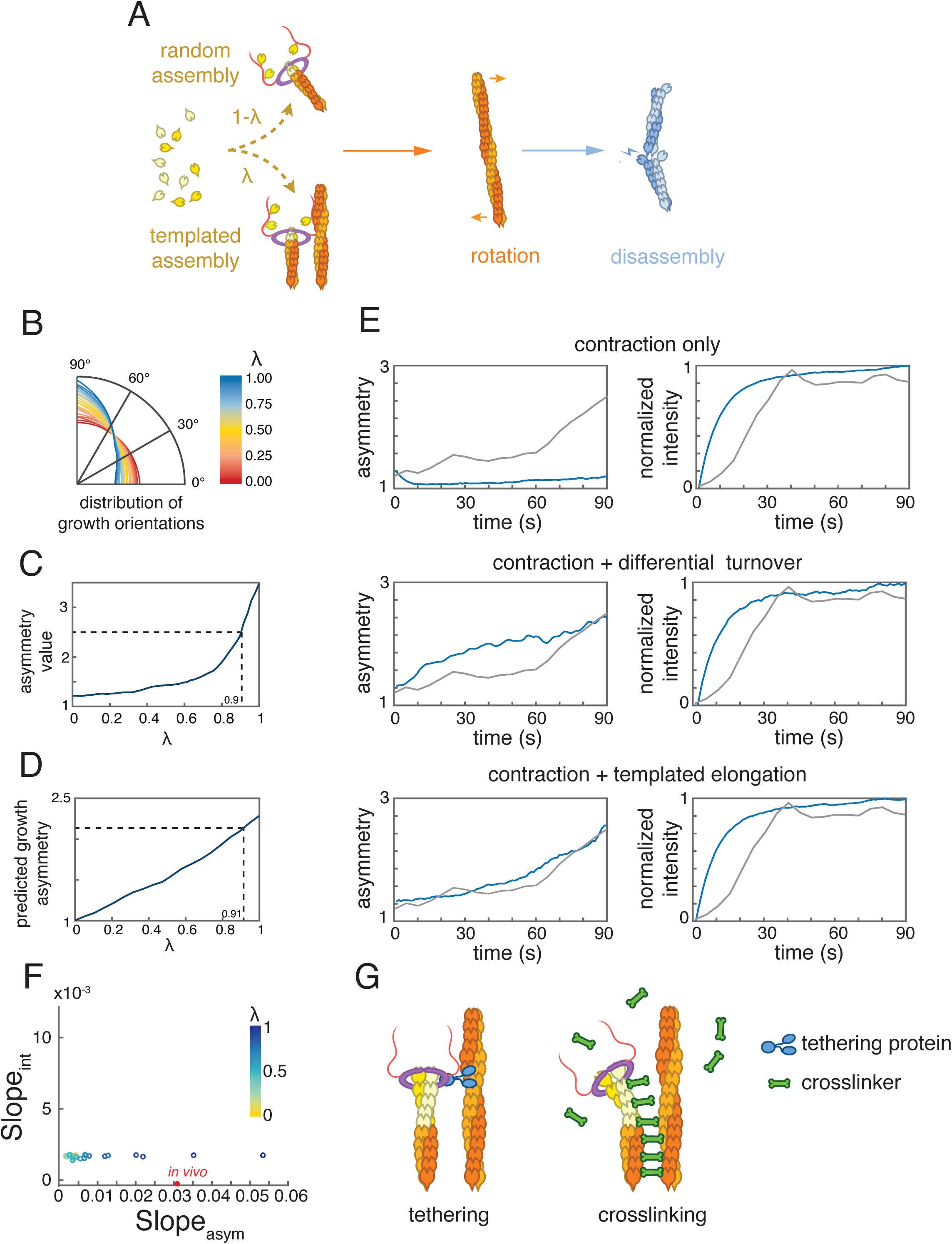
Filament-guided filament assembly (FGFA) increases the effective lifetime of filament orientation, and is sufficiently strong to explain the rapid emergence of filament alignment, given measured contraction and disassembly rate. (A) A simple model for filament-guided filament assembly: a fraction (*λ*) of growing filaments use existing filaments as guides to orient their growth. (B) The predicted distributions of filament orientations after 90 seconds, for different values of *λ*, given measured contraction and filament disassembly rate (Figure 3E and I respectively). (C) Predicted asymmetry values for the distributions in (B). Dashed line indicates the value of *λ* required to produce the measured asymmetry value in Figure 1E (*λ*=0.9). (D) Predicted filament growth asymmetry for different values of *λ*, given the distribution of filament orientations measured in late Phase II (Figure 1E). Dashed line indicates the value of *λ* required to produce the measured asymmetry value in Figure 4G (*λ*=0.91). (E) A comparison of model predictions for three different scenarios. For “contraction only” (top), we assumed constant assembly and disassembly at the measured rate (Figure 3I). For “contraction plus orientation-dependent disassembly” (middle), we assumed constant assembly and orientation-dependent disassembly. We selected parameter values (k_min_ = 0.0301, k_max_ = 0.1501, K = 80, Figure S4A) for which the mean turnover and final asymmetry values matched the measured values, and for which the slope of normalized F-actin density and asymmetry value vs time during Phase II best-matched the measured values (see Figure S4C, see Experimental Procedures for details). For “contraction plus filament-guided filament assembly” (bottom), we used the measured disassembly rate and chose *λ* = 0.9 to match the final asymmetry value measured *in vivo*. In each case, model predictions (blue curves) are compared to the measured values (grey curves). For all three scenarios, we used the measured contraction rate (Figure 3E). (F) Values of asymmetry slope vs. intensity slope predicted for different values of *λ* in the filament-guided filament assembly model, given measured contraction and disassembly rates. Red circle indicates the value measured *in vivo*. (G) Schematic view of two possible mechanisms for filament-guided filament assembly (FGFA). Left: Formin is tethered to an existing filament by unknown tethering proteins during Formin-mediated filament elongation. Right: newly assembled filaments are rapidly crosslinked to an existing filaments by crosslinkers.

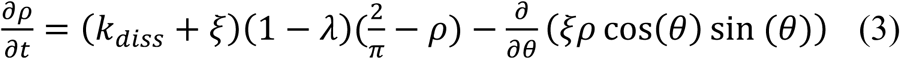

Comparing Equations (2) and (3), we see that the effect of filament-guided filament assembly is to scale the time for relaxation of filament orientations by a factor 1 − *λ*, yielding an effective relaxation time 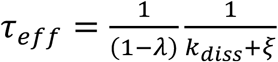.

We then returned to simulations to ask how high must *λ* be to achieve the measured asymmetry of equatorial filaments after 90 seconds, given measured contraction and filament disassembly rates. We found that simulations reproduced the measured asymmetry for *λ* = 0.9 (Figure 6B-C). Moreover, compared to simulations based on equatorial contraction with constant turnover, or equatorial contraction with orientation-dependent disassembly (using best-fit parameters), simulations that combined FGFA (for *λ* = 0.9) with measured rates of equatorial contraction and filament disassembly rate correctly captured the time-dependent changes in filament density and asymmetry observed *in vivo*, with a fast initial rise in density to a stable plateau, and an accelerating rise in asymmetry value (Figure 6E). In essence, FGFA allows separate control over the lifetimes of individual filaments and the effective lifetime of filament orientation (Figure S4 F). This in turn allows for an initially fast rise in filament density governed by the balance of filament assembly/disassembly, followed by later sharp rise in filament asymmetry driven by compressive flow.

Finally, to estimate the value of *λ in vivo*, we used simulations to ask how large would *λ* have to be, given the measured asymmetry of equatorial filament orientations (Figure 1F), to produce the distribution of equatorial CYK-1 movement directions observed *in vivo* (Figure 4G). This analysis yielded an estimated value for *λ* of 0.91, which is comparable to the value required in simulations. We conclude that FGFA could be sufficient to explain the emergence of equatorial filament alignment despite the rapid turnover of individual filaments.

## Discussion

A classical model for cytokinesis in animal cells propose how compressive cortical flows, driven by equatorial activation and/or polar inhibition of cortical contractility, could concentrate and align actin filaments at the equator during contractile ring assembly (Reymann et al., 2016; Salbreux et al., 2009; Spira et al., 2017; White and Borisy, 1983). For this mechanism to produce a given degree of filament alignment, the rate of equatorial cortex compression must be sufficiently high, and/or the local memory of filament alignment must be sufficiently long. Recent work has confirmed that equatorial contraction rates are sufficiently fast to produce in principle the degree of filament alignment observed during contractile ring assembly in *C. elegans* zygotes (Khaliullin et al., 2018; Reymann et al., 2016), but only if the memory of filament alignment is greater than 2-3 minutes. Here, we have used single molecule analysis to show that during contractile ring assembly, polymerized actin has a mean lifetime of ∼ 8 seconds, which is far too short for individual filaments to encode a stable memory of network alignment. Moreover, comparing the output of simple models to measurements of filament density, alignment and turnover over time, we find that the rapid emergence of filament alignment cannot be explained by preferential stabilization of correctly oriented filaments. Any such mechanism predicts a progressive increase in filament lifetime and density as filaments become more favorably aligned, which we do not observe.

Instead, our direct observations of filament assembly *in vivo* identify a mechanism by which the zygote encodes a long-term memory of filament alignment, in which filaments assembled by Formin/CYK-1 use existing filaments as guides to orient their growth. Support for this mechanism comes from three observations: First, we find that filament growth orientations correlate strongly and locally with existing filament orientations in both wild type and myosin-depleted embryos. Second, two-color imaging of filament growth trajectories in relation to existing filaments shows that newly assembling filaments have a strong tendency to elongate along paths defined by existing filaments, and to turn when they encounter existing filaments/bundles. Third, observations of filaments that grow into the equatorial region reveals a very strong tendency for filaments to reorient and align their growth with the equatorial axis, a tendency that increases with increasing alignment of equatorial filaments. Thus, existing filaments act as local templates to continuously influence the alignment of newly elongating filaments, a property that we refer to as filament-guided filament assembly (FGFA).

What is the molecular basis for FGFA? One possible scenario is that one or more factors tether a growing filament’s barbed end to the side of an existing filament. In principle, this tethering could be mediated by the simultaneous association of CYK-1 dimers with a growing filament’s barbed-end and adjacent filaments (Figure 6F, left), as recently reported for another elongation factor Ena/VASP (Harker et al., 2019). However, Ena/VASP forms tetramers that can associate with multiple filaments while promoting processive filament elongation (Harker et al., 2019). In contrast, Formin dimers use both FH2 domains to associate with the barbed-end of the elongating actin filament to maintain rapid processive elongation. Thus additional modes of Formin binding to F-actin would be required to tether a processively elongating filament to an existing one.

A second, and perhaps more plausible scenario, is that one or more crosslinking proteins act to “zipper” growing filaments with existing filaments (Figure 6F, right), allowing the former to inherit the orientation of the latter. Two possible candidates in early *C. elegans* embryos are the crosslinkers PLST-1/Plastin (Fimbrin) (Ding et al., 2017) and ANI-1/Anillin (Descovich et al., 2018; Maddox et al., 2005; 2007). Both PLST-1 and ANI-1 accumulate in the contractile ring during cytokinesis (Ding et al., 2017; Maddox et al., 2007), and can bundle F-actin in vitro (Ding et al., 2017; Tian et al., 2015). *plst-1* mutant zygotes display variably penetrant cytokinesis defects, ranging from delayed constriction to a complete failure of cytokinesis (Ding et al., 2017), while strong depletion of ANI-1 abolishes asymmetric furrow ingression (Maddox et al., 2007). In both cases, these defects are associated with reduced and/or delayed alignment of filaments within the contractile ring (Ding et al., 2017; Reymann et al., 2016). However, reduced alignment could also be caused by reduced recruitment of equatorial myosin II and/or reduced cortical connectivity (Ding et al., 2017; Reymann et al., 2016; Tse et al., 2011), leading to reduced cortical flows, rather than being caused by changes in the orientation of filament growth. Thus, further experiments are required to test specific roles for these proteins in FGFA.

Regardless of the underlying molecular details, FGFA provides a powerful mechanism for growing filaments to inherit a memory of previous filament’s orientations. The strength of this effect will depend on the frequency with which a growing filament encounters existing filaments/bundles and the probability per encounter that a filament reorients its growth. This strength can be characterized by the average fraction of time *λ* that new filaments grow with orientations given by existing filaments. Our simple theoretical analysis shows that FGFA increases the effective local lifetime of filament orientation by a factor 1/(1-*λ*) over the lifetime of individual filaments. Thus in principle, by increasing the efficiency of FGFA (i.e. by driving *λ* -> 1), it is possible to create an arbitrarily long-lived memory of local filament alignment. Our stochastic simulations, constrained by measured equatorial contraction rates and mean filament lifetimes, predict that *λ* must be greater than 0.9 to explain the degree of filament alignment observed during contractile ring assembly. By comparing the orientations of equatorial filaments, and equatorial filament growth (Figure 6D), we estimate that *λ* ∼ 0.9. However, this is likely to be an underestimate, because reorientation of filament growth is progressive over the growth trajectories of individual filaments (Figure 5E) and particle tracking errors and photobleaching of CYK-1::GFP bias our observations to the beginnings of growth trajectories. Thus, structural memory of filament alignment conferred by FGFA makes a dominant contribution to building filament alignment during contractile ring assembly.

That said, additional forms of structural memory may also contribute to contractile ring assembly. For example, myosin minfilaments turn over more slowly than single actin filaments (Carvalho et al., 2009) and may also be aligned by flow (Singh et al., 2019). ANI-1/Anillin may locally crosslink and/or stabilize F-actin and Myosin II to promote their asymmetric accumulation within the contractile ring and asymmetric furrow ingression (Maddox et al., 2007; Tian et al., 2015). Local curvature induced by ring constriction may also feedback to enhance local filament alignment {Dorn:2016cc}. In principle, any of these mechanisms could synergize with FGFA to enhance filament alignment during cytokinesis.

Why might cells rely on FGFA to build a structural memory of filament alignment, rather than increasing the lifetimes of single filaments? Rapid filament turnover may be important for local network homeostasis – i.e. to maintain uniform filament density and prevent local tearing and/or clumping of contractile ring components (Chew et al., 2017). At the same time, filament turnover provides an effective way to dissipate local resistance to the rapid remodeling of actin networks that underlies flow and filament realignment (McFadden et al., 2017). Indeed, in a network of cross-linked filaments undergoing axial compression, filaments cannot realign without some degree of local filament buckling or inter-filament sliding (Bidone et al., 2017; Murrell et al., 2015). Thus, FGFA provides a way to maintain a selective memory of network deformation, while maintaining filament homeostasis and allowing rapid dissipation of local resistance to network deformation.

Although we have focused here on the initial phase of contractile ring assembly, FGFA is also likely to make an essential contribution to maintaining filament alignment during ring constriction and furrow ingression. For compressive flow to align actin filaments, it must be anisotropic (White and Borisy, 1983). The axial compressive flows that accompany ring assembly satisfy this requirement, but during later cytokinesis, the equatorial cortex compresses both axially and circumferentially as it enters the furrow (Khaliullin et al., 2018). Rapid ∼isotropic compression may help to concentrate Myosin II to maintain a constant rate of ring constriction through cytokinesis (Carvalho et al., 2009; Khaliullin et al., 2018), but it will no longer contribute to building filament alignment. Therefore, mechanisms that maintain local filament alignment are likely to be even more important during ring constriction. Importantly, if each encounter of a growing filament with an existing filament/bundle carries a fixed probability of realigning filament growth, then the strength of FGFA will increase with filament density and degree of alignment and thus it will be strongest during late stages of ring assembly and during ring constriction.

Compressive flows underlie the formation of transient dynamically aligned filament arrays in many other functional contexts, including cell motility, immune cell synapse formation, wound healing and tissue morphogenesis (Burnette et al., 2011; Hotulainen and Lappalainen, 2006; Mandato and Bement, 2001; Murugesan et al., 2016; Sehring et al., 2014). In all of these contexts, biology must solve a similar problem – how to build and maintain filament alignment in the face of continuous turnover of individual filaments. Even more stable structures such as stress fibers or junctional actin belts undergo steady filament turnover and cells must therefore implement strategies to ensure that newly added filaments are correctly aligned. It will be interesting to see to what extent filament-guided filament assembly provides a common solution to this widespread challenge.

## Supplementary Figure Legends

**Figure S1.**
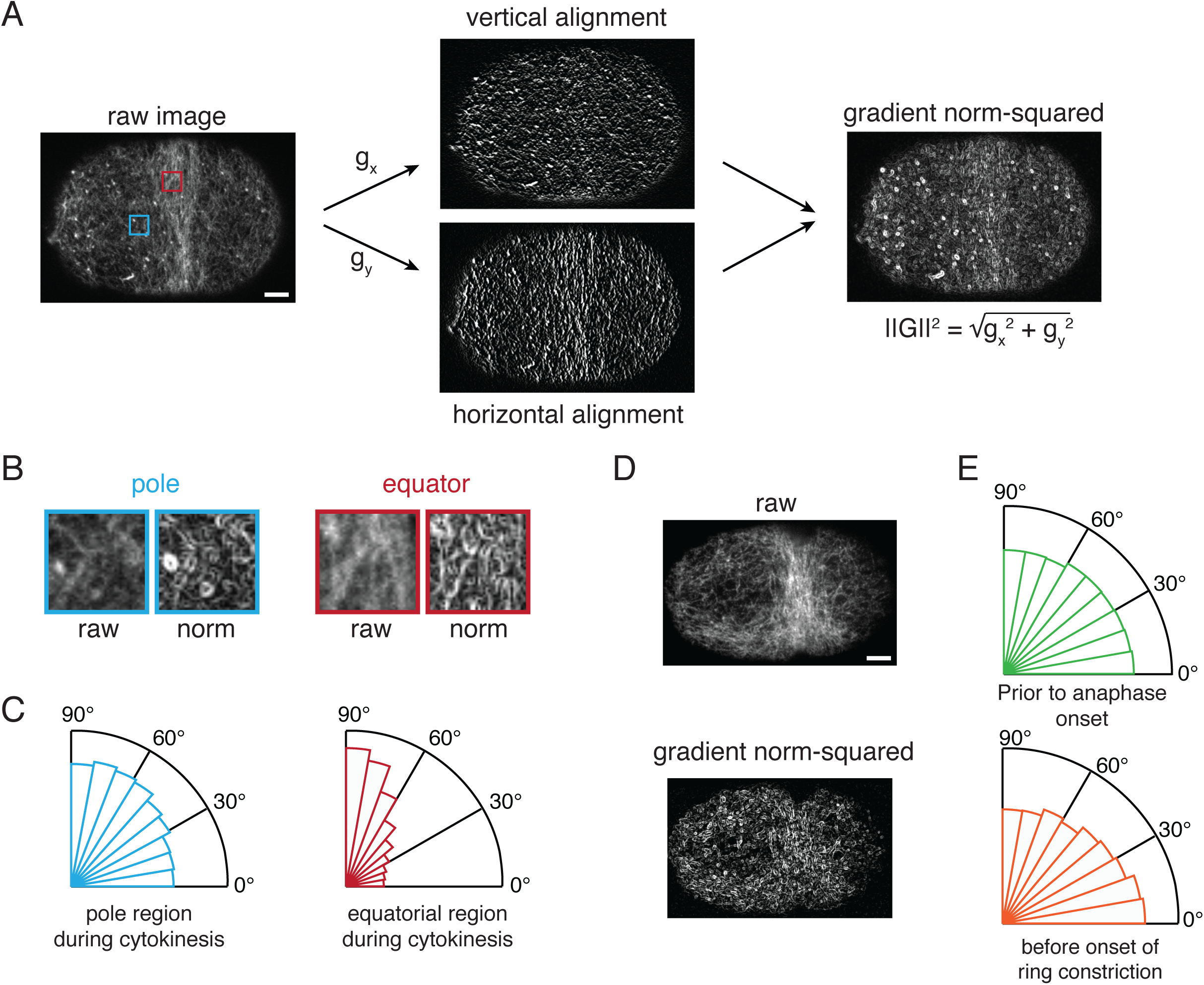
(related to Figure 1). Measurement of actin filament alignment. (A) (Left) Raw image of a fixed phalloidin-stained embryo. (Middle top and bottom) The same image subjected to Sobel filters 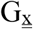 (top) and 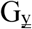 (bottom) to highlight horizontal and vertical gradients of fluorescence intensity. (Right) Gradient norm-squared image in which pixel intensity is proportional to the squared norm of G = (G_x_, G_y_). Scale bar = 5 µm. (B) Magnified views of the regions indicated by colored boxes in (A) comparing raw signal and gradient norm squared. size = 4×4 µm. (C) Weighted distribution of polar and equatorial filament orientations averaged over n = 6 fixed phalloidin-stained embryos, that were fixed in cytokinesis, before the onset of rapid furrow ingression. (D) Comparison of raw and gradient norm-squared images for a live embryo expressing GFP::UTR just before the onset of rapid furrow ingression. Scale bars = 5 µm. (E) Weighted distribution of polar filament orientations (anterior pole) in live embryos expressing GFP_UTR just before anaphase onset (left) and just before the onset of rapid furrow ingression (right), averaged over n = 5 embryos.

**Figure S2.**
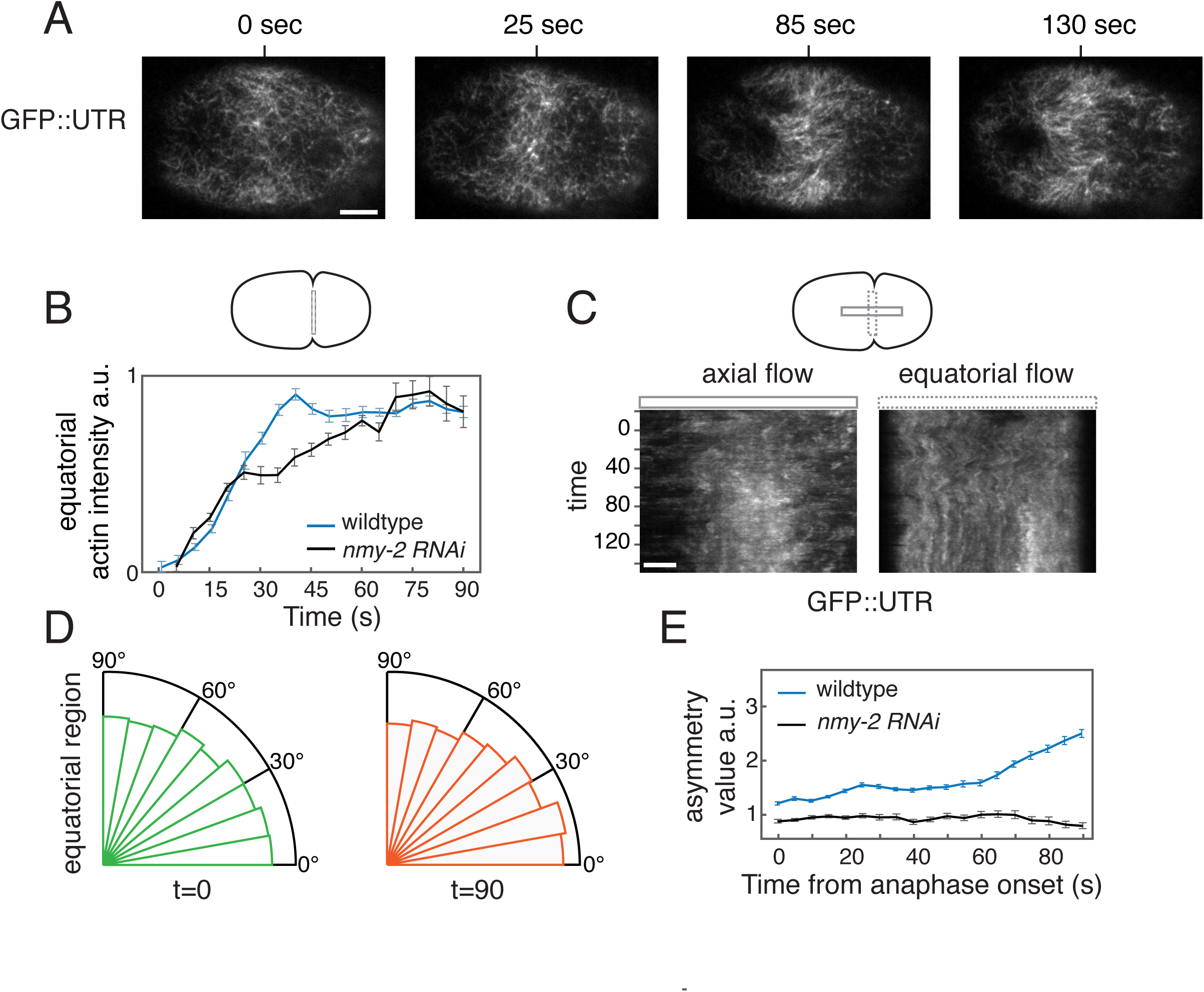
(related to Figure 1&4). Analysis of filament orientation in myosin-depleted embryos. (A) Sequence of near-TIRF images from anembryo expressing GFP::UTR and strongly depleted of myosin II by *nmy-2 (RNAi)*. Times measured relative to anaphase onset. Scale bar = 5 µm. (B) Plot of mean equatorial actin filament intensity vs time in (n = 5) *nmy-2(RNAi)* embryos. Top schematic indicates the regions used for measurements of probe densities (box). Wild type data from Figure 1B is shown for comparison. Error bars indicate SEM. (C) Kymographs showing axial (left) and equatorial (rotational) (right) cortical flows in the embryo shown in (A). Top schematic indicates the regions used to make axial (solid box) and equatorial (dashed box) kymographs. Scale bar = 3 µm. (D) Radial histograms showing the distribution of local filament orientations at the equator at anaphase onset (left, t=0) and 90 seconds later (n=5 embryos). (E) Plot of mean asymmetry value vs time in (n = 5) *nmy-2(RNAi)* embryos. Data for wild type embryos from Figure 1F is shown for comparison. Error bars indicate SEM.

**Figure S3.**
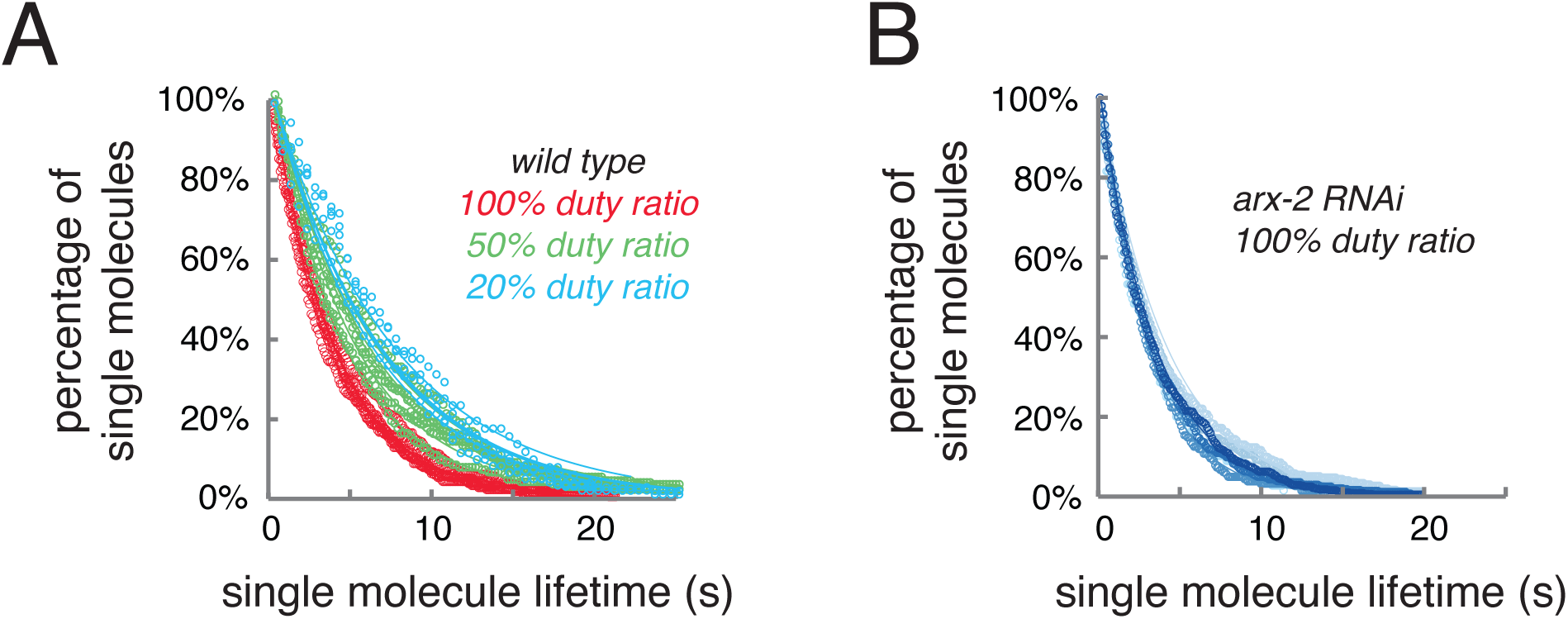
(Related to Figure 3). Single molecule lifetime measurements at different duty ratios in wild type and *arx-2(RNAi)* embryos. (A) Loss curves plotting the fraction of single molecule trajectories with a given lifetime recorded at the equator during cytokinesis under illumination at three different duty ratios. Loss rates inferred from exponential fits to these data are shown in Figure 3H. (B) Loss curves plotting the fraction of single molecule trajectories with a given lifetime recorded at the equator during cytokinesis in *arx-2(RNAi)* embryos using 100% duty ratio.

**Figure S4.**
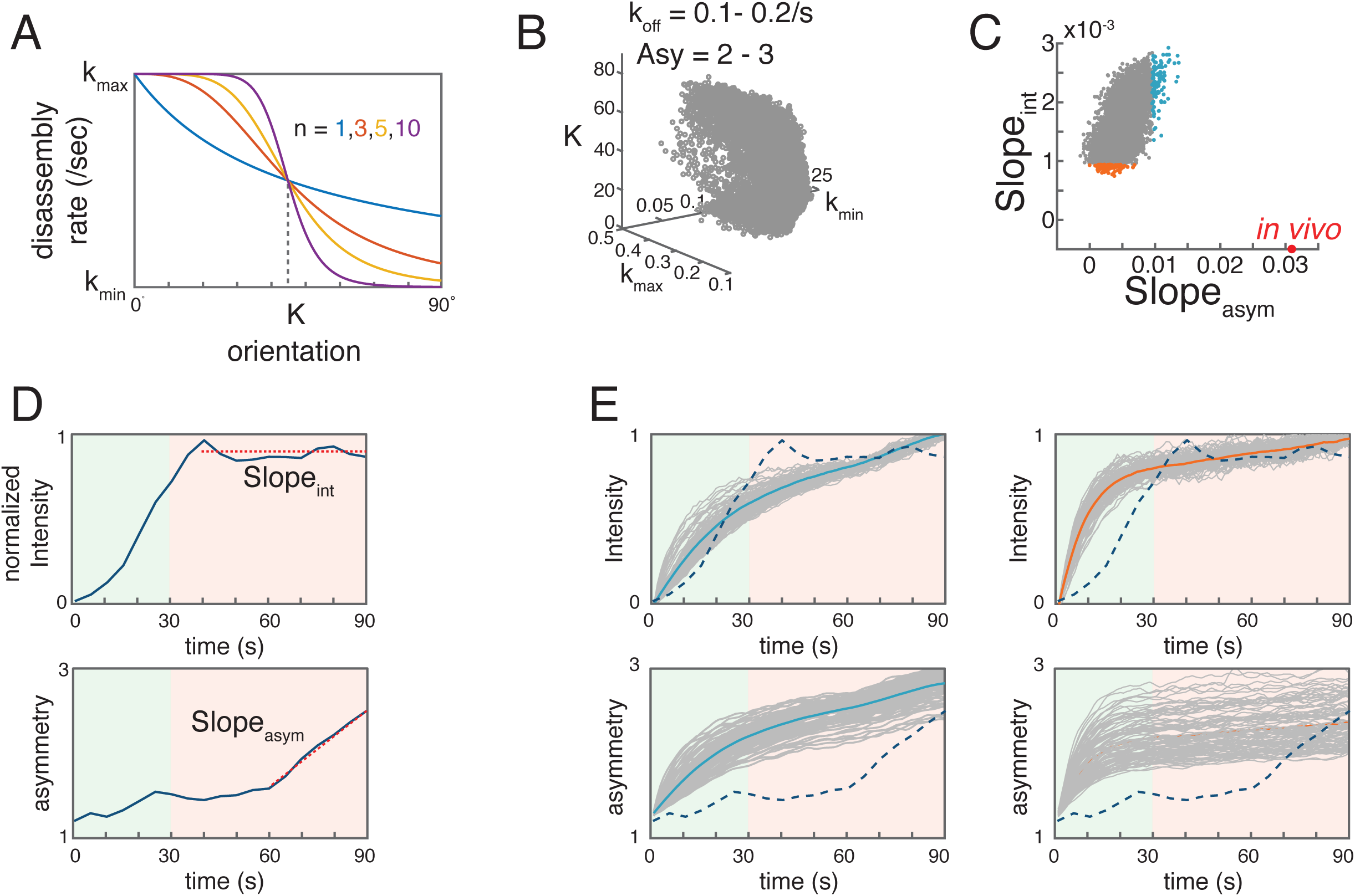
(Related to Figure 3) Preferential stabilization of oriented filaments cannot explain the emergence of filament alignment. (A) Using a Hill function to represent the dependence of disassembly rate on filament orientation. 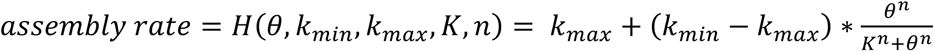. 0° is axial, 90° is equatorial. (B) Scatter plot showing the set of values for k_min_, k_max_, K, and n for which simulations satisfy two criteria: mean disassembly rate for the last 30 seconds is between 0.1/s – 0.2/s, and the asymmetry value after 90s is between 2 and 3. Parameter values were sampled with the ranges: k_max_ ∈ (0, 0.5]; k_min_ ∈ (0, 0.2]; k ∈ (0, 90]; n ∈ [1, 10]. (C) Scatter plot of asymmetry slope vs. intensity slope (defined in (D)) for all sets of parameters from (B). Red circle marks the measured *in vivo* values. Blue circles mark the 100 simulations whose asymmetry slope value best matches *in vivo* asymmetry slope. Orange circles mark the 100 simulations whose intensity slope value best matches *in vivo* intensity slope. (D) Illustration of the method used to measure the intensity slope (top) and asymmetry slope (bottom). Dashed lines indicate the time interval in which the slope was measured by fitting a line to the data. Dark blue curves show the *in vivo* measurements from Figure 1B&F. (E) Plots of intensity (top) and asymmetry value (bottom) vs time for the parameter sets indicated by filled circles in (C). Grey curves show the results for individual parameter sets. Colored solid curves show the average of all grey curves. Dark blue dash lines show the *in vivo* measurements from Figure 1B&F.

## Supplementary Movie Legends

**Movie S1 (related to Figure 1) Two-color imaging of F-actin and Myosin II dynamics during cytokinesis**. Near-TIRF movie of a one-cell *C. elegans* embryo during cytokinesis expressing UTR::GFP to label F-actin, and endogenously tagged NMY-2::mKate2. Top panel: UTR::GFP; Middle panel: NMY-2 :mKate2; bottom panel: merged UTR::GFP (green) and NMY-2::mKate2 (red). Scale bar = 5 µm. Time compression is 30:1.

**Movie S2 (related to Figure 3) Single molecule imaging of Actin::GFP to monitor cortical flow and filament turnover during cytokinesis**. Near-TIRF movie of a one-cell *C. elegans* embryo during cytokinesis expressing Actin::GFP at single molecule levels. Top panel: raw movie. Bottom panel: the raw movie overlaid with all single molecule trajectories (labeled in red circles) of length >= 2 frames. Scale bar = 5 µm. Time compression is 18:1.

**Movie S3 (related to Figure 4) Single particle imaging of CYK-1::GFP to monitor actin filament growth during cytokinesis**. Near-TIRF movie of a one-cell *C. elegans* embryo near anaphase onset expressing endogenously-tagged CYK-1::GFP. Top panel: raw movie. Middle panel: 13-frame moving minimum intensity projection highlighting the stationary fraction of CYK-1 particles. Bottom panel: Difference between raw and minimum intensity projection highlighting the mobile fraction of CYK-1 particles (see Experimental Procedures for details). Scale bar = 5 µm. Time compression is 5:1.

**Movie S4 (related to Figure 4) Filtering CYK-1 trajectories to identify fast directionally moving CYK-1 particles**. Near-TIRF movie of a one-cell *C. elegans* embryo near anaphase onset expressing endogenously-tagged CYK-1::GFP, with subsets of tracked particles overlaid as red circles (see Experimental Procedures for details). Top panel: all particle trajectories of length >= 2 frames. Second panel: Subset of trajectories with length >= 10 frames. Third panel: Subset of trajectories with length >= 10 frames, and directional movement. Bottom panel: Subset of trajectories with length >= 10 frames, directional movement, and 0.8 <= mean speed <= 2.5 µm/sec. Scale bar = 5 µm. Time compression is 5:1.

**Movie S5 (related to Figure 4) CYK-1::GFP trajectories represent growing filaments**. Zoomed view of the anterior cortex of a one-cell *C. elegans* embryo near anaphase onset expressing endogenously-tagged CYK-1::GFP (red) and lifeAct::mCherry (green). The embryo was partially depleted of Myosin-2 to suppress cortical rotation (see Experimental Procedures for details). Scale bar = 2 *μm*. Time compression is 9:1.

**Movie S6 (related to Figure 5) Growing filaments turn as they encounter existing filaments/bundles**. Zoomed view of the cortex of a one-cell *C. elegans* embryo near anaphase onset expressing endogenously-tagged CYK-1::GFP (magenta) and lifeAct::mCherry (cyan). The embryo was partially depleted of Myosin-2 to suppress cortical rotation (see Experimental Procedures for details). Top, middle and bottom panels highlight three different examples of a fast-moving CYK-1::GFP particle, marking a growing filament, which turns as it encounters an existing filament or bundle. Left and right panels show images processed to enhance the Actin and CYK-1 signal (see Experimental Procedures for details). Yellow circle in right panels indicates the CYK-1 particle of interest. Scale bar = 2 *μm*. Time compression is 9:1.

**Movie S7 (related to Figure 5) Growing filaments turn as they enter the equatorial region**. Near-TIRF movie of examples of CYK-1 particles traveling into the equatorial region. The white lines mark equatorial region, and the red circles highlight the CYK-1 particle of interest. Scale bar = 3 *μm*. Time compression is 5:1

## Experimental Procedures

### *C. elegans* Culture and Strains

We cultured *C. elegans* strains under standard conditions (Brenner, 1974). See Supplemental Experimental Procedures for a list of mutations and transgenes used in this study. Unless otherwise specified, strains were provided by the Caenorhabditis Genetics Center, which is funded by the National Center for Research Resources.

### RNA interference

We performed RNAi using the feeding method (Timmons et al., 2001). Unless otherwise specified, bacteria targeting specific genes were obtained from the library of (Kamath et al., 2003). L4 larvae were transferred to feeding plates and then cultured at various temperatures for various times before imaging: 24-30 hours at 20 °C for *nmy-2(RNAi)* for strong NMY-2 depletion (Figure 4H&S2), 12-16 hours at 20 °C for *nmy-2(RNAi)* for mild NMY-2 depletion (Figure 4D&5A), 16-24 hours at 24 °C for *arx-2(RNAi)* (Figure 3I&S3B). For experiments involving *nmy-2 (RNAi)*, we verified strong loss of function by complete failure of first cleavage, and mild loss of function by lack of cortical rotation. For experiments involving *arx-2(RNAi)*, we verified strong loss of function by extreme anterior displacement of the pseudocleavage furrow (Shivas and Skop, 2012).

### Live imaging

For all live imaging experiments, we mounted embryos in egg salts containing ∼100 uniformly sized polystyrene beads (15.6 mm diameter; Bangs Laboratories, #NT29N) to achieve mild compression of the embryo surface that is suitable for single molecule imaging and particle-tracking analysis, while maintaining a uniform degree of compression across experiments (Robin et al., 2014).

For two-color imaging of F-actin (GFP::UTR) and myosin (NMY-2::mKate2) or CYK-1 (CYK-1::GFP and F-actin (Lifeact::mCherry), we used a Nikon Ti-E inverted microscope equipped with solid state 50mW 481 and 561 Sapphire lasers (Coherent), a TIRF illuminator, and a Ti-ND6-PFS Perfect Focus unit. A laser merge module (Spectral Applied Research; LMM5) equipped with an acousto-optical tunable filter (AOTF) allowed rapid (1-2 msec) switching between excitation wavelengths. We collected near TIRF images using a CFI Apo 1.45 NA oil immersion TIRF objective, with 1.5X magnification, onto an Andor iXon3 897 EMCCD camera, yielding a pixel size of ∼107nm. Image acquisition was controlled by Metamorph software.

For single molecule/single particle tracking analysis of GFP::Actin or CYK-1::GFP, we used an Olympus IX50 inverted microscope equipped with an Olympus OMAC two-color TIRF illumination system, a CRISP autofocus module (Applied Scientific Instrumentation), and a 1.45 NA oil immersion TIRF objective. Laser illumination at 488 nm from a 50-mW solid-state Sapphire laser (Coherent) was delivered by fiber optics to the TIRF illuminator. Images were magnified by 1.6x and collected on an Andor iXon3 897 EMCCD camera, yielding a pixel size of 100 nm. Image acquisition was controlled by Andor IQ software.

For all experiments, we set the laser illumination angle to a standard value that was chosen empirically to approximately maximize signal-to-noise ratio while maintaining approximately even illumination across the field of view.

### Single-particle detection and tracking

We performed single-particle detection and localization using a MATLAB implementation (http://people.umass.edu/kilfoil/downloads.html) of the Crocker and Grier method (Crocker, 1996; Pelletier et al., 2009). Briefly, in each image, the method uses a band pass filter to highlight roughly circular regions below a characteristic size (the feature size), in which the pixel intensity exceeds the background. The regions in which the maximum intensity exceeds a user-defined threshold are identified, and their centroids are determined to sub-pixel resolution as the center of mass of pixel intensity within a pixelated circular mask centered on the original maximum. We used a feature size of 3 and chose thresholds subjectively to optimize detection for different types of particles (Actin::GFP and CYK-1::GFP).

We performed particle-tracking analysis using freely available µTrack software (Jaqaman et al., 2008, https://github.com/DanuserLab/u-track). µTrack first links particles frame to frame and then links these short segments into longer sequences. Both linking steps use statistical models for particle motion to compute costs for different possible linkage assignments (particle appearance, disappearance, displacement, fusion, and fission) and then identify the assignments that globally minimize these costs. For all analyses reported here, we used a motion model provided with µTrack that represents a mixture of Brownian and directed motion. We allowed the possibility of “gaps” in trajectories due to transient failure to detect particles in individual frames. For each embryo, we overlaid the raw movie with tracked particles to verify tracking accuracy, and we chos parameters for particle detection and tracking (thresholds and length of gaps) that minimize tracking errors. We previously verified the accuracy of these for measuring actin filament turnover (Robin et al., 2014). For our analyses of CYK-1 or Actin particle movements, all of our experiments were performed at particle densities for which the major tracking errors are failures to link together real particle trajectories, and these errors will have negligible effects on measurements of particle speed and direction.

### Measuring filament density and orientation

For analysis of filament density and orientation (Figure 1), we imaged embryos expressing UTR::GFP and NMY-2::mKate2, using stream acquisition mode with 30% laser power, and 100 msec exposure times, for both GFP and RFP channels. At each time point, and for each channel, we collected a stack of 3 focal planes, in 0.1 um increments, starting from the cortical surface and going inward. Then we used an average intensity projection to obtain a mean intensity for each pixel.

To monitor changes in the densities of equatorial F-actin and Myosin II, we measured the total intensity of UTR::GFP and NMY-2::mKate2 signals within the equatorial ROI over time (5 pixels x 122 pixels).

To estimate the distribution of filament orientations within equatorial ROIs (6 *μm* in width. From hereafter 6 *μm* equatorial region is used to measure equatorial actin filament disassembly rate, the orientation of the elongation of equatorial filament, and the boundary to decide whether a CYK-1 particle moves into the equatorial region or not.), we used the Sobel operator to identify local gradients of fluorescence intensity, which are sharp in directions orthogonal to individual filaments/bundles. We convolved raw images with the Sobel operator defined by 3×3 kernels 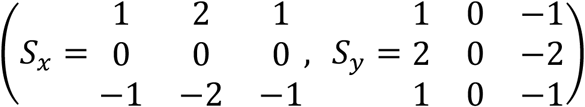 to compute the x and y components *G*_*x*_ and *G*_*y*_ of the local fluorescence intensity gradient (Figure S1A). For each pixel, 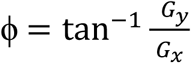 is the gradient’s direction, and 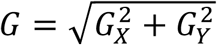 its magnitude. Therefore, we took the positive angle orthogonal to ϕ, 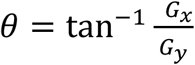, to be a local estimate of filament orientation, with magnitude G. To estimate the distribution of filament orientations within a given ROI, we assigned each pixel within that ROI to the appropriate orientation bin (0 <= *θ* <= 90°), weighted by the G, to emphasize the sharp local gradients associated with individual filaments/bundles.

### Measuring filament asymmetry values

To estimate a single scalar asymmetry value from distributions of filament orientations, we measured the density of filament orientations between 80° - 90° (*ρ* _80−90_) and 0° - 10° (*ρ* _0−10_), and defined the orientation asymmetry to be the ratio of *ρ* _80−90_ :: *ρ* _0−10_.

### Measuring filament disassembly rate and axial contraction rates

To image single molecules of Actin::GFP for measurements of cortical flow, we collected data in stream acquisition mode using 30% laser power for both wild type embryos and *arx-2(RNAi)* embryos). For axial contraction rate, we collected data using 450 msec exposure times. For disassembly rate, we collected data using 100 msec exposure times. We performed particle detection and tracking as described above, and then performed all subsequent analyses using custom scripts written in Matlab and R (available upon request).

To measure axial contraction rate, in MatLab we selected the subset of trajectories with lifetimes greater than 5 frames to exclude false positives. We confirmed the reliability of tracking by overlaying the resulting trajectories on the original image data. We applied a linear transformation to map positional coordinates onto X and Y axes aligned with the embryo’s AP axis. Finally, we calculated the frame-to-frame displacement for each point of each trajectory and exported these data into R.

In R we took the x-axial component of the frame-to-frame displacements over all trajectories and binned them with respect to time (bin size = 30 frames = 13.5 seconds) and with respect to axial position (bin size = 10 pixels = 1 um). For each time bin, we performed a Loess fit (https://www.rdocumentation.org/packages/stats/versions/3.6.2/topics/loess) to estimate the mean axial velocity as a function of axial position. We then computed the forward time difference of mean velocity for each axial position to approximate the derivative of axial velocity (axial strain rate) at each point.

To measure cortical disassembly rates, we first selected all trajectories beginning within a proscribed region (equatorial or polar) and window of time (30 seconds before the onset of ring constriction). We then aligned the beginnings of all single molecule trajectories to construct a standard decay curve plotting the percentage of trajectories that remain after an interval of time *τ* vs t. These decay curves were well-fit by single exponentials, yielding an estimate of the single molecule disappearance rate, which is the sum of the F-actin disassembly rate and the rate of single molecule photobleaching.

To estimate single molecule photobleaching rates at 30% laser power, we measured decay curves for many individual embryos, holding laser power constant at 30% while varying the duty ratio (fraction of time the laser is on) and then fit the resulting data to *k*_*disappearance*_= *k*_*disassembly*_ + *dr* * *k*_*photobleach*_ to estimate *k*_*disassembly*_ and *k*_*photobleach*_.

### Measuring actin filament assembly using CYK-1::GFP

To monitor Formin-dependent filament assembly during cytokinesis, we imaged embryos expressing GFP-tagged CYK-1 (CYK-1::GFP) expressed form either the endogenous locus (Padmanabhan et al., 2017) or as an integrated transgene {MiMi:2012ka}. For both strains, and for both wild type and *nmy-2(RNAi)* embryos, we collected data in stream acquisition mode using 100% laser power and 61 msec exposure. In raw movies, the majority of CYK-1::GFP signal appeared as diffraction-limited speckles, and we could detect two general classes of speckles - stationary and moving. To focus our analysis on the moving speckles, we computed a moving minimum intensity projection of the raw data, with a 13-frame window, to highlight the stationary fraction of CYK-1::GFP speckles. We then subtracted this stationary fraction from the original image data to construct a sequence of images in which the moving particles could be more readily detected and tracked.

We performed particle detection and tracking on these processed data, as described above (see Movie S4, all tracked). We then performed several additional filtering steps to select for trajectories representing fast directional movement characteristic of formin-mediated filament elongation. First, we manually selected a subset of longer trajectories, which corresponded by eye to rapidly moving CYK-1 particles. For this subset, we divided each trajectory into smaller fragments (10 frames = 0.3 seconds per fragment), and computed a distribution of average mean speeds to serve as a reference for selecting shorter trajectories.

Next, we considered the subset of all trajectories with lengths greater than or equal to 10 frames (see Movie S4, Length >= 10). We decomposed each of these selected trajectories into shorter 10-frame segments. Then we used a method previously developed by Jaqaman and colleagues (Jaqaman et al., 2008) to select for directional movement. This method assigns an asymmetry parameter (S) to each trajectory based on the eigenvalues *λ*_1_ and *λ*_2_ of its variance-covariance matrix.

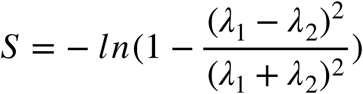

Based on simulated data, Jaqaman et al (2008) found that the S value required to achieve > 90% detection accuracy decreases with increasing trajectory length, approaching a plateau at S ∼ 1.5 as the trajectory length increases above 10 frames. Therefore, we selected for further analysis the subset of 10-frame trajectory segments with S > 1.5 (see Movie S4, Length >= 10, directional). Finally, we selected the subset of these segments with an average mean speed within the range of the means measured for the subset of manually chosen long trajectories (see Movie S4, Length >= 10, directional, 0.8 <= mean speed <= 2.5). We plotted the selected trajectory segments over the original data to confirm accuracy of particle tracking and trajectory segment selection. Finally, we estimated the movement direction for each selected trajectory segment from the positional difference between the first and last frames of the segment.

### Visualizing CYK-1 movements along actin filament bundles

To visualize movements of CYK-1 speckles along individual filament bindles, we imaged embryos expressing transgenic CYK-1::GFP and Lifeact::mCherry, mildly depleted of myosin II to abolish acute rapid equatorial cortical rotation during cytokinesis. We used alternating 50 msec exposures for each channel. To enhance visualization of actin filaments, we applied a moving average of 5 frames to the Lifeact::mCherry data. To enhance visualization of moving CYK-1 speckles, we first applied a gaussian blur of 1.3 to the CYK-1::GFP data, then a bandpass filter to suppress pixel noise and spatial variations at wavelengths greater than the characteristic particle size. Then we subtracted a moving minimum average of 3 frames to suppress signals associated with stationary CYK-1 speckles and bleed through from Lifeact::mCherry signal. Finally, we applied a gamma filter of 1.5 to highlight the bright CYK-1::GFP speckles.

### Analyzing CYK-1 trajectories moving into the equatorial region

To capture CYK-1 trajectories moving into the equatorial region, we imaged embryos expressing transgenic CYK-1::GFP {MiMi:2012ka}, which is expressed at lower levels than the endogenous protein. We manually selected 202 CYK-1::GFP particles from 13 embryos that crossed a boundary into the equatorial region, and manually tracked each particle. We aligned all trajectories with respect to their point of entry into the equatorial region and by reflecting trajectories about the AP and/or equatorial axis to place them all into the same quadrant. To calculate the distribution of the movement direction of those trajectories, we divided each trajectory into 5 frame segments, and for each segment, we estimated the movement direction from the positional difference between the first and last frames.

### Modeling Procedures

We constructed a simple model that predicts how the distribution of filament orientations at the equatorial cortex evolves through a combination of local assembly, reorientation by contractile flow, and disassembly. We considered a patch of equatorial cortex, with width W and fixed height H, which contracts at a constant strain rate *ξ*. We denoted the density of filaments with orientation *θ* by *ρ*(*θ, t*). We assumed that filaments assemble at a rate *k*_*ass*_(*θ*) and disassemble at a rate *k*_*diss*_(*θ*). We assumed further that the cortical filament network undergoes a locally affine deformation, such that filaments within this network rotate at a rate 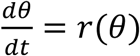. Finally, we imposed a moving boundary condition - the left and right boundaries move as the patch deforms to satisfy 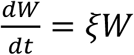, such that there is no flux of filaments across the left or right boundaries, and the number of filaments in the domain changes only through assembly and disassembly. Given these assumptions, we wrote an equation that describes how the distribution of filament orientations within the patch evolves over time:

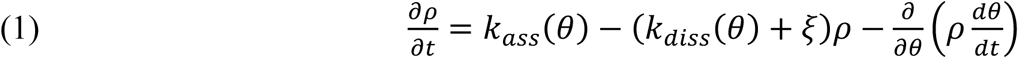

To derive an expression for 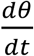, we focused attention on a single filament within the network, with orientation *θ*, and with axial and circumferential length components *L*_*x*_ and *L*_*y*_ respectively. Affine deformation of the network changes *L*_*x*_, but leaves *L*_*y*_ unchanged. Orientation is related to the axial and circumferential lengths *L*_*x*_ and *L*_*y*_ by 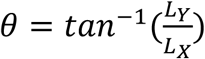

Taking a time derivative, applying the chain rule, and using 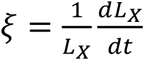 we obtained:

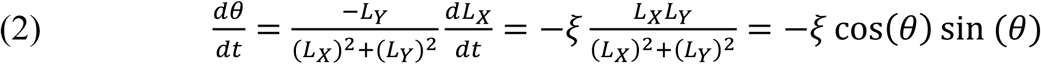

### Orientation-independent filament assembly and disassembly

We initially considered the simplest scenario, in which *k*_*ass*_ and *k*_*diss*_ are independent of filament orientation. Thus:

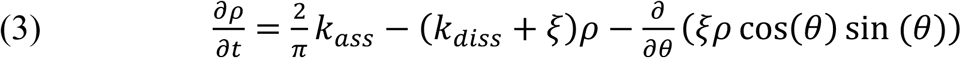

Where the factor 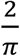 has been chosen so that the total filament assembly rate is *k*_*ass*_. Scaling *ρ* in Equation (3) by 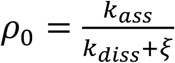 yields:

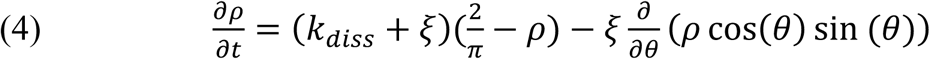

From this it can be seen that the time evolution of the distribution of filament orientations depends only on the value of *k*_*diss*_ and *ξ*. The second term in Equation (4) represents the buildup of alignment due to flow, while the first term represents the relaxation of the filament orientations towards an isotropic distribution with relaxation time 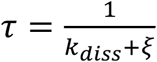.

### Orientation-independent filament assembly and disassembly

To study how filament guided filament assembly affects the evolution of filament orientation under compressive flow, we considered a scenario in which a fraction of filaments (w) elongate using an existing filament as a guide, while the rest (1-w) assemble with random orientation. Accordingly, we rewrote Equation (3) as follows:

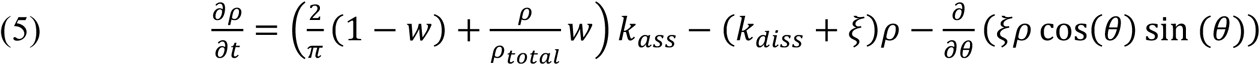

Where 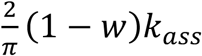 is the rate of randomly oriented assembly and 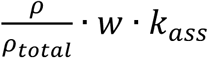 is the rate of filament-guided assembly. Note that we scaled these rates by 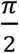 and 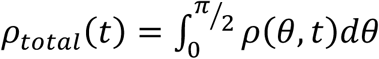 so that the total rates of randomly-oriented and filament-guided assembly are (1 − *w*) *k*_*ass*_ and *w* · *k*_*ass*_ respectively. The dynamics of *ρ*_*total*_(*t*) are given by:

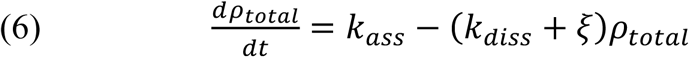

As above, we scaled *ρ* by 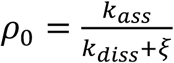 to obtain:

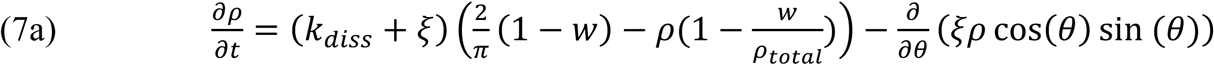

and

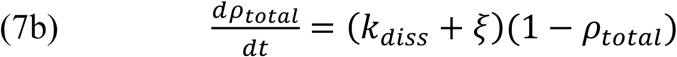

Equation (7b) implies that *ρ*_*total*_ → 1, for times 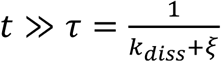, thus at long times, Equation (7a) can be approximated by:

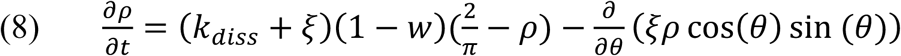

Comparing Equations (4) and (8), we see that the effect of filament guided filament assembly is to scale the time for relaxation of filament orientations by a factor 1 − *w*, yielding an effective relaxation time

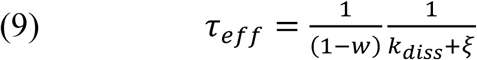

In particular, when all filament assembly is guided by existing filaments (*w* = 1), *τ* _*eff*_ → *∞*, and the time to build filament alignment is set only by the contraction rate.

